# Heparin-derived theranostic nanoprobes overcome the blood brain barrier and target glioma in murine model

**DOI:** 10.1101/2022.01.07.475195

**Authors:** Sumanta Samanta, Vadim Le Joncour, Olivia Wegrzyniak, Vignesh Kumar Rangasami, Harri Ali-Löytty, Taehun Hong, Ram Kumar Selvaraju, Ola Aberg, Jons Hilborn, Pirjo Laakkonen, Oommen P. Varghese, Olof Eriksson, Horacio Cabral, Oommen P. Oommen

**Affiliations:** Bioengineering and Nanomedicine Lab, Faculty of Medicine and Health Technology, Tampere University, 33720 Tampere, Finland; Translational Cancer Medicine Research Program, Faculty of Medicine, University of Helsinki, Helsinki, Finland; Science for Life Laboratory, Department of Medicinal Chemistry, Uppsala University, Uppsala, Sweden; Surface Science Group, Photonics Laboratory, Tampere University, P.O. Box 692, FI-33014 Tampere, Finland; Department of Bioengineering, Graduate School of Engineering, The University of Tokyo, 7-3-1 Hongo, Bunkyo-ku, Tokyo 113-8656, Japan; Polymer Chemistry Division, Department of Chemistry, Ångström Laboratory, Uppsala University, 75121 Uppsala, Sweden

**Keywords:** heparin, nanoparticles, blood-brain-barrier, glioma, radioimaging

## Abstract

The poor permeability of theranostic agents across the blood-brain-barrier (BBB) significantly hampers the development of new treatment modalities for neurological diseases. We have discovered a new biomimetic nanocarrier using heparin (HP) that effectively passes the BBB and targets glioblastoma. Specifically, we designed HP coated gold nanoparticles (HP-AuNPs) that were labeled with three different imaging modalities namely, fluorescein (FITC-HP-AuNP), radioisotope ^68^Gallium (^68^Ga-HP-AuNPs), and MRI active gadolinium (Gd-HP-AuNPs). The systemic infusion of FITC-HP-AuNPs in three different mouse strains (C57BL/6JRj, FVB, and NMRI-nude) displayed excellent penetration and revealed uniform distribution of fluorescent particles in the brain parenchyma (69-86%) with some accumulation in neurons (8-18%) and microglia (4-10%). Tail-vein administration of radiolabeled ^68^Ga-HP-AuNPs in healthy rats also showed ^68^Ga-HP-AuNP inside the brain parenchyma and in areas containing cerebrospinal fluid, such as the lateral ventricles, the cerebellum, and brain stem. Finally, tail-vein administration of Gd-HP-AuNPs (that display ∼3 fold higher relaxivity than that of commercial Gd-DTPA) in an orthotopic glioblastoma (U87MG xenograft) model in nude mice demonstrated enrichment of T1-contrast at the intracranial tumor with a gradual increase in the contrast in the tumor region between 1h-3h. We believe, our finding offers the untapped potential of HP-derived-NPs to deliver cargo molecules for treating neurological disorders.

---

Despite enormous progress in the field of bionanomaterials for drug delivery applications, there is limited success in designing nanocarriers that could effectively deliver therapeutic cargo molecules and imaging modalities to the intact brain. This is attributed to the blood-brain barrier (BBB) that plays a major role in protecting the central nervous system from toxins and pathogens. The unique selectivity of the BBB also limits the delivery of therapeutic drug molecules and thereby curtails our ability to treat malignancies such as neurological disorders after early diagnosis.^[1]^ As a result, several promising drugs fail to achieve therapeutic efficacy and therefore neurological diseases such as Alzheimer’s disease, Parkinson’s disease, multiple sclerosis, brain tumor, etc. remain difficult to treat. Discovering new biomimetic pathways that can deliver drug payloads and imaging modalities efficiently through the BBB and to targets in the brain is therefore of paramount importance.^[2,3]^

Currently, the delivery of therapeutic molecules across BBB can be characterized into three major pathways. The first strategy involves the binding of drug molecules, nanoparticles (NPs), or surfactants to serum proteins, such as albumin or ApoE, that interact with the BBB receptors or transporters to allow entry to the brain parenchyma. For example, poly(L-lactide-co-glycolide) (PLGA)-derived nanoparticles^[4]^ and surfactant polysorbate 80 (or tween 80) PLGA nanoparticles invade BBB by binding to ApoE after infusion in the blood.^[5]^ The second strategy involves receptor targeting transcytosis where a ligand is installed on the NP surface to interact directly with the BBB receptors, such as the transferrin receptor,^[6,7]^ glucose receptor,^[8]^ integrin receptor,^[9–11]^ or by conjugating NPs with cell-penetrating peptides.^[12]^ The third strategy for paracellular delivery of drugs across BBB involves transient fenestration of the BBB’s endothelium using small molecules such as mannitol^[13]^ or recombinant proteins, such as human vascular endothelial growth factor^[14]^ or targeting tight junction proteins such as claudin-5.^[15]^ The transient disruption of the BBB brings its complexity and risk the infiltration of immune cells and other undesired biomolecules to the brain. Despite all these developments, we have limited success in treating patients with neurological diseases,^[16]^ suggesting an unmet need to develop promising delivery tools that are safe and have the potential to treat diverse brain diseases.

Heparin is a natural glycosaminoglycan that displays anti-coagulant properties and is known to bind growth factors, show antiangiogenic characteristics, and suppress inflammation.^[17]^ Earlier studies have reported that unfractionated heparin is effective in suppressing neuroinflammation in the experimental autoimmune encephalomyelitis (EAE) model of multiple sclerosis, albeit, at low doses.^[18]^ However, no further studies are reported, presumably due to the dose-limiting effects. Since unfractionated heparin (UHP) has limited ability to penetrate the BBB,^[19]^ the ultralow-molecular weight heparin^[19]^ or heparin-derived oligosaccharides C3, were developed for efficient BBB permeability.^[20]^ The low molecular weight heparin (LMWH or enoxaparin) and heparin-derived oligosaccharide(s) (HDO) are clinically used for the management of neurological disorders, such as stroke and Alzheimer’s disease.^[21,22]^ Moreover, as 20%–50% of patients with metastatic brain tumors develop venous thromboembolism, LMWH is currently used in the clinic to treat cancer-related thrombotic complications.^[23]^ Administration of LMWH to patients with a primary brain tumor or secondary metastatic brain tumor is found to be beneficial to mitigate venous thromboses^[24]^ with an acceptable risk of bleeding.^[25]^

Several groups have also developed heparin-derived nanocarriers for drug delivery applications.^[26]^ Heparin coating of nanoparticles provides stealth properties and enhances the bioavailability as such a coating lowers the risk of elimination by the reticuloendothelial system in the liver, and spleen and prevents adsorption by serum proteins.^[27]^ Heparin coated liposomes were also found to display ≈1.5-fold higher stability than PEGylated liposomes, resulting in a higher accumulation of particles in tumor tissue in murine melanoma (B16F10) model.^[27]^ Recently, heparin-derived nanocarrier decorated with integrin-binding RGD peptides were designed for targeting glioma in a mouse orthotopic model^[28]^ by exploiting the unique property of the RGD peptides to promote BBB permeability.^[9,11]^ Since there is sufficient literature that provides convincing evidence of the BBB permeability of LMWH and HDOs, we hypothesized that heparin-based nanoparticles could potentially mimic LMWH and breach the BBB, which could be used for the delivery of cargo molecules. Since heparin is a natural biopolymer with well-established bioactivity, biodegradability, and immune response, we believe heparin-derived nanocarriers could potentially be a powerful arsenal for delivering cargo molecules to the brain tissue without producing any toxic metabolites.

## Results and discussion

To verify our hypothesis of BBB permeability of HP, we synthesized the fluorescently-tagged UHP by conjugating fluorescein thiosemicarbazide (FTSC) to heparin (HP-FTSC) following a carbodiimide coupling strategy that was previously optimized in our laboratory.^[29]^ As a negative control, we developed FTSC conjugated chondroitin sulfate polymer (CS-FTSC). Conjugation of hydrophobic FTSC to the biopolymers resulted in the formation of self-assembled micelles of 150 and 170 nm hydrodynamic size for HP and CS respectively, as determined by the dynamic light scattering (DLS) experiments (Figure S1 in Supporting information or SI). The degree of FTSC conjugation in HP and CS was estimated to be 3.21±0.9% and 2.2±0.5% respectively using the FTSC extinction coefficient of 78 000 M^−1^ cm^−1^ at 492 nm. However, the ^1^H NMR analysis of HP-FTSC and CS-FTSC in D_2_O (Figure S1C and Figure S1D) did not display the aromatic signals of FTSC suggesting a core-shell self-assembly.^[30,31]^

### *In-vitro* evaluation of HP-FTSC permeability across artificial BBB

In order to screen and evaluate the potential of HP-FTSC to cross the BBB, we used the animal-free *in-vitro* BBB model in a dish. This model, based on co-cultures of either human or mouse endothelial cells and astrocytes, on the two sides of a transwell membrane, accurately predicts the permeability of the compounds and small molecules across the BBB.^[32–35]^ We compared the potential penetration of CS-FTSC and HP-FTSC nanoparticles (40 µg/mL, 100 µg total) to the passively diffusing, fluorescein dye after 48 h. Permeability calculation revealed the significantly reduced ability of the CS-FTSC to cross the *in-vitro* BBB, however, the HP-FTSC were abundantly detected in the brain side (astrocyte layer) of the insert (Figure 2A). We then verified the distribution of the fluorescence at the cellular level, on the *in-vitro* BBB using confocal microscopy taken through the endothelial and astrocyte monolayers of the *in-vitro* BBB model. After 48 h, both the human (Figure 2B) and the murine (Figure 2C) *in-vitro* BBB showed evidence of increased amounts of HP-FTSC on the brain/astrocyte side, whereas the CS-FTSC was accumulating on the blood/endothelial side with reduced amounts detected on the brain/ astrocyte side compartment. We then examined whether the facilitated passage of HP-FTSC would be due to the potential cytotoxic effect on the endothelial/astrocyte cells. Unhealthy endothelial/astrocyte cocultures lead to interruptions of the tight junction cellular wall and endothelial fenestrations resulting in increased permeability. Using the MTT cell viability assay,^[36]^ we incubated the different cells used for establishing the BBB, e.g. human endothelial cells (HuAR2T) (Figure 2D), murine endothelial cells (bEND3) (Figure 2F), normal human astrocytes (NHA) (Figure 2E), and immortalized murine astrocytes (HIFkoAs) (Figure 2F) with the same concentration of HP-FTSC (40 µg/mL) of for 48 h. As a positive control for cytotoxicity, we used the microtubule stabilizer doxorubicin (1 µM) under the same conditions. When compared to untreated cells, doxorubicin significantly decreased the viability of all tested cells, while treatment with HP-FTSC nanoparticles did not greatly alter the cell viability (over 95% viability). Taken together, these *in-vitro* results indicate that HP-FTSC constructs can cross the BBB without toxic side effects for the neurovascular unit.

**Figure 1.**
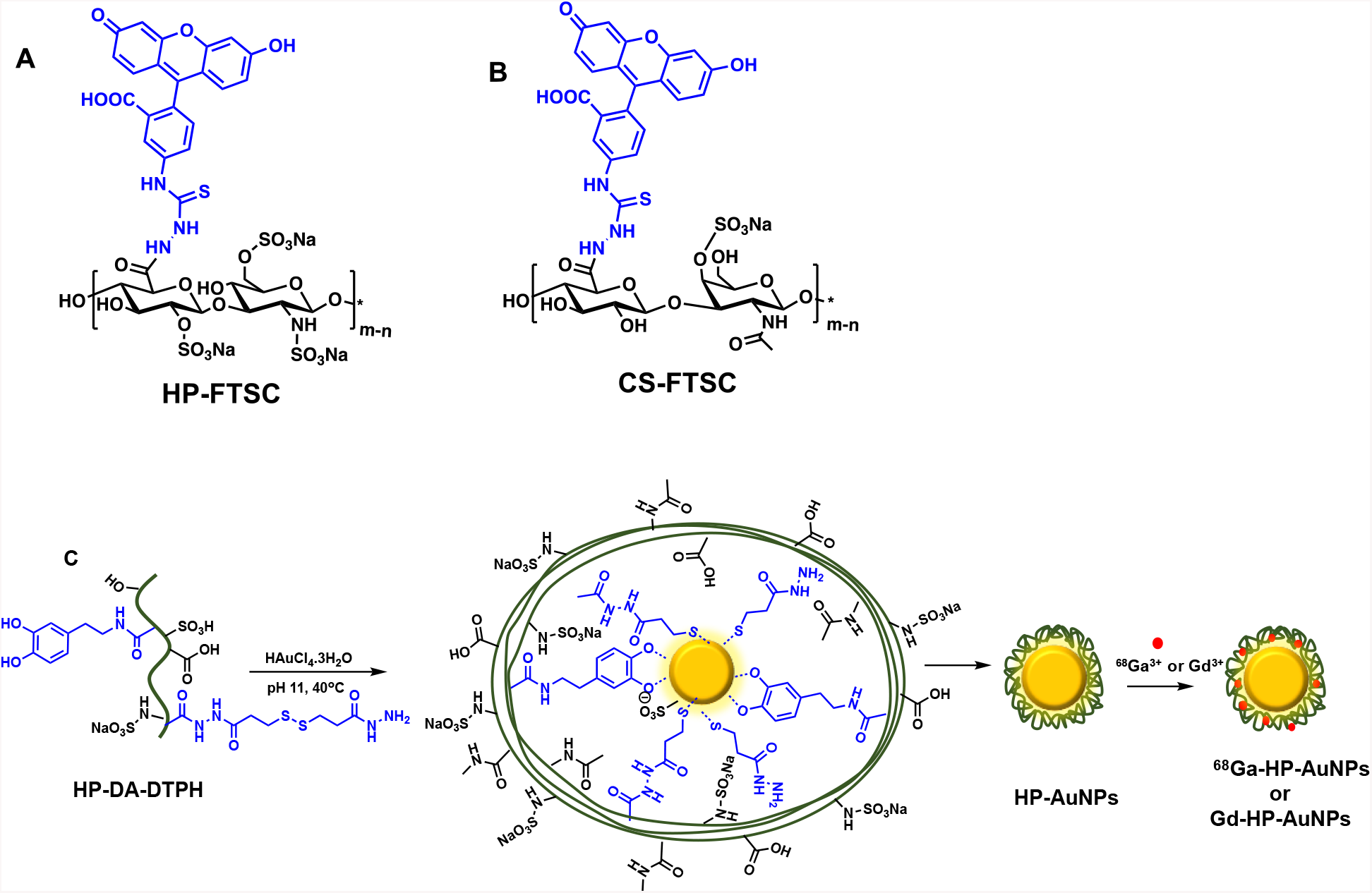
Chemical structure of (A) HP-FTSC, (B) CS-FTSC, (C) Schematic representation of the synthesis of HP-AuNPs. HP modified with dopamine and DTPH reduces gold chloride to spherical gold nanoparticles and the anionic functional groups on the nanoparticle surface efficiently complex metallic imaging agents.

**Figure 2.**
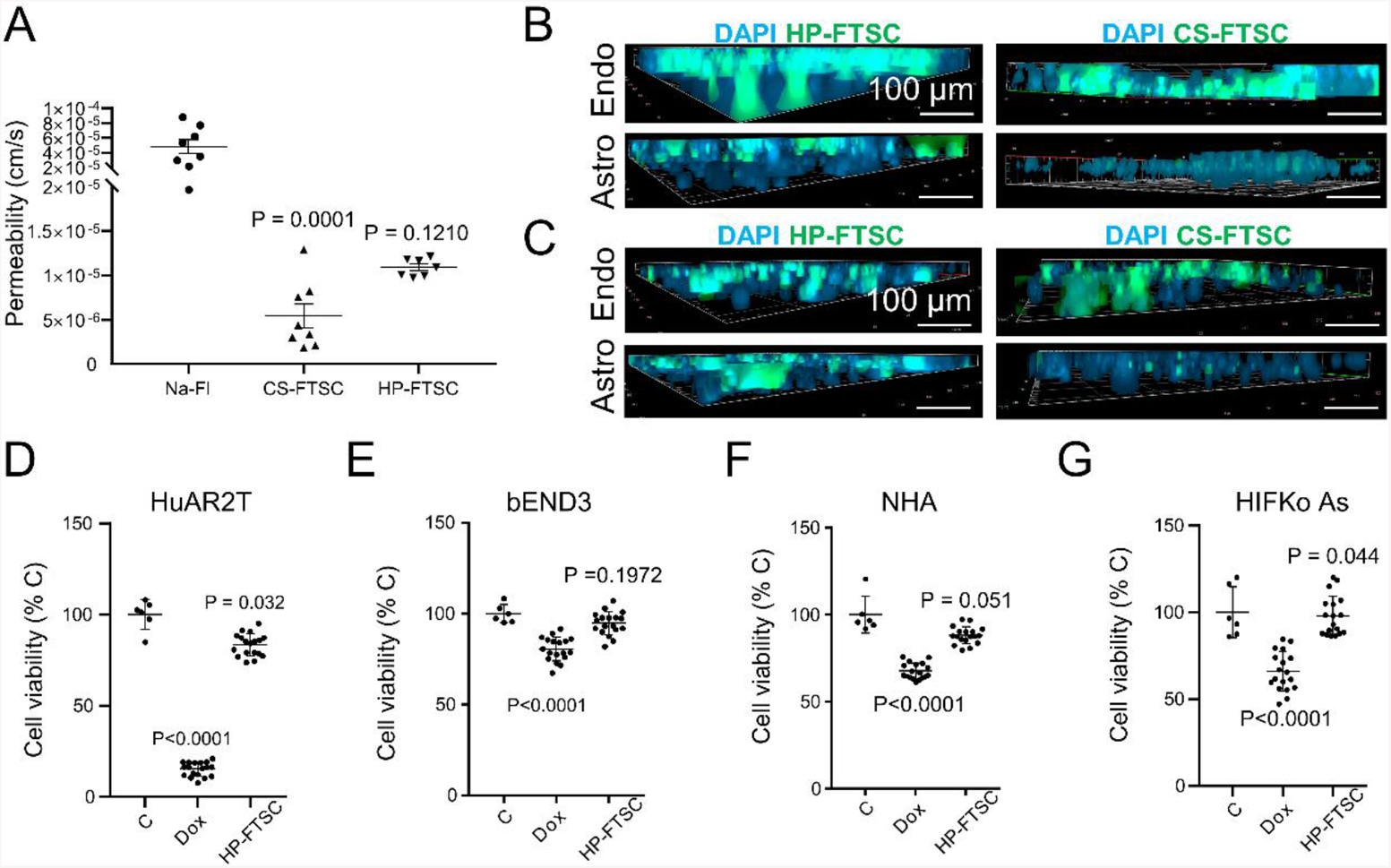
Assessment of the BBB permeability of HP-FTSC and CS-FTSC 48 h post-incubation. **A** Calculation of the passage through the *in-vitro* BBB of the positive control molecule (sodium-fluorescein, Na-Fl) and comparison with CS and HP nanoparticles (100 µg, each). Lower permeability values indicate retention of the compound on the blood/endothelial side of the chamber and decreased BBB penetration. **B**, Tridimensional z-stack confocal micrographs of the blood/endothelial (top) and brain/astrocyte (bottom) sides of the human *in-vitro* BBB inserts. HP-FTSC (left) and CS-FTSC (right) are conjugated to fluorescent FTSC (green) and cell nuclei are counterstained with DAPI. **C**, Tridimensional z-stack confocal micrographs of the blood/endothelial (top) and brain/astrocyte (bottom) sides of the murine *in-vitro* BBB inserts. HP-FTSC (left) and CS-FTSC (right) (green) and cell nuclei are counterstained with DAPI. **D**, Cell viability of the HuAR2T endothelial cells 48 h after treatment with cytotoxic drug doxorubicin (Dox), HP-FTSC, or untreated control cells (C). **E**, Cell viability of the bEND3 endothelial cells 48 h after treatment with cytotoxic (Dox), HP-FTSC, or untreated control cells (C). **F**, Cell viability of the NHA astrocyte cells 48 h after treatment with cytotoxic Dox, HP-FTSC, or untreated control cells (C). **G**, Cell viability of the HIFko astrocyte cells 48 h after treatment with Dox, HP-FTSC, or untreated control cells (C). P-values were calculated using the 2-way ANOVA with Dunnett’s multiple comparison test to the control.

### HP-FTSC cross the blood-brain-barrier *in vivo* and are detected in the cerebral cortex

In order to validate the data obtained from the *in-vitro* BBB experiments, adult male FVB mice were infused with 4 mg/kg HP-FTSC or CS-FTSC (100 µg in 200 µl of PBS) through the caudal vein. Control animals were injected with 200 µl of vehicle (PBS). Brain tissues were collected after 1 h (n = 3) and 24 h (n = 3) in order to verify the BBB penetration and brain distribution. We then compared their respective green fluorescent signal emitted from the FTSC conjugate in 30 µm thick mouse brain sections by confocal microscopy. Interestingly, i. v. injection of HP-FTSC demonstrated high levels of fluorescence (green) signal in the brain parenchyma of the cerebral cortex (DAPI blue), outside the blood vessel capillaries (red) 1 h post-injection (Figure 3A), which were still detectable in the brain after 24 h of infusion, suggesting relatively good stability and slow clearance of the HP-FTSC from the brain parenchyma. Corroborating with the *in-vitro* results, evaluation of CS-FTSC demonstrated negligible fluorescence signal related to the intraparenchymal diffusion of the CS construct (green) in the brain cerebral cortex (DAPI, blue), after 1 h (Figure 3B). This signal was not observable after 24 h post-injection in the cerebral cortex.

**Figure 3.**
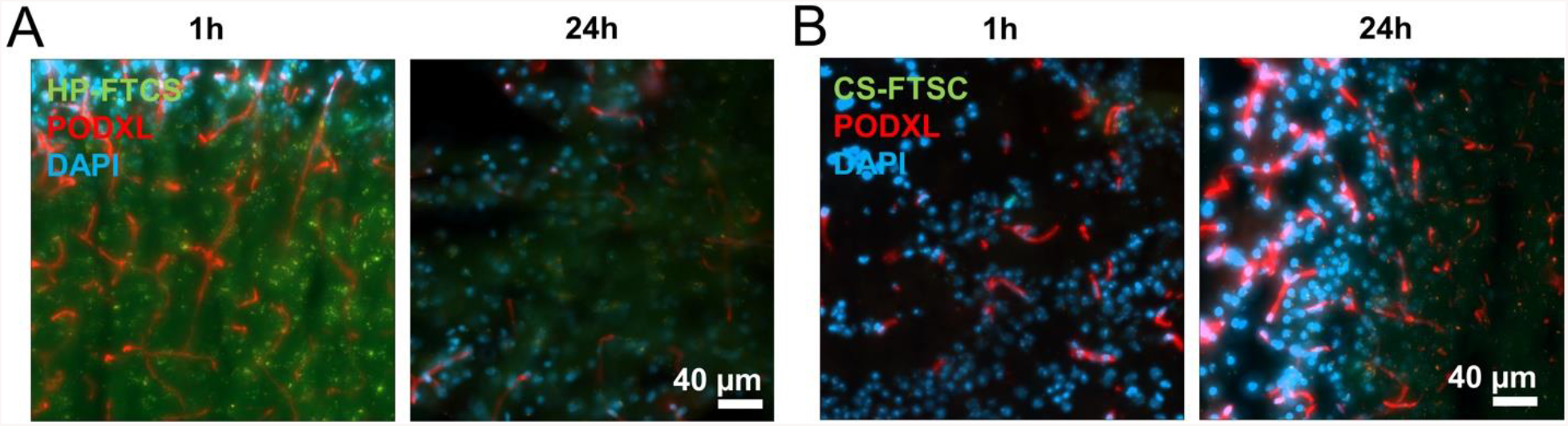
Intracerebral detection of HP-FTSC and CS-FTSC in 1 h and 24 h post intravenous injection. Brain micrographs of a male FVB mouse injected with (**A)** HP-FTSC and **(B)** CS-FTSC (4 mg/kg, 100 µg in 200 µL, green) through the caudal vein. After 1 h or 24 h, the brain from each group was collected and immunofluorescent labeling of the endothelial podocalyxin (PODXL, red) and DAPI nuclear counterstaining (blue) were performed.

### Synthesis and characterization of HP coated gold nanoparticles (HP-AuNPs)

In our endeavor to identify new biomimetic nanocarriers that could deliver model drug molecules and imaging agents to the brain, we synthesized heparin-coated gold nanoparticles (HP-AuNPs). The HP-AuNPs were synthesized following a green chemistry approach using our modified protocol that we developed for synthesizing chondroitin sulfate-based nanocarriers.^[29]^ To achieve this, we first grafted disulfide hydrazide groups and dopamine moiety to the unfractionated heparin (UHP) by conjugating 3,3’-dithiobis(propanoic hydrazide) or DTPH (8% with respect to the carboxylate groups) and dopamine units (13% with respect to the carboxylate groups) following carbodiimide coupling chemistry as estimated by UV-Vis spectroscopy. The NMR analysis of HP and its derivatives are complex^[37]^ as HP polymer is composed of *N*-acetyl glucosamine and glucuronic acid as the repeat units with dispersed iduronic acid in its backbone, thus making the assignment of reference signal difficult. In addition, the complex sulfation pattern in the HP further complicates the NMR analysis. However, the ^1^H-NMR spectra further confirmed the successful conjugation of dopamine moiety onto the HP backbone (Figure S2B in SI), which was estimated to be 7.7% with respect to the *N*-acetyl signal from the N-acetyl glucosamine moiety. Both the DTPH units^[29]^ and the dopamine moiety^[38]^ are capable of reducing Au^III^ to Au^0^ forming gold nanoparticles. Thus, HP functionalized with dopamine and DTPH would generate stable gold nanoparticles with good polydispersibility and stability following green chemistry, without using any organic solvents or reducing agents. With the optimized ratio of HAuCl_4_ and heparin-Dopa-DTPH conjugate, we obtained gold nanoparticles (HP-AuNPs) with the hydrodynamic size of ≈86 nm as confirmed by DLS measurement, which displayed size of 50-60 nm gold nanoparticle core with 10 nm HP corona as evidenced by transmission electron microscopy (Figure S4 in SI). The free hydrazide moiety derived from the DTPH on the AuNP surface was used to conjugate fluorescein by reacting the AuNPs with fluorescein isothiocyanate (FITC) that renders stable covalent conjugate by the formation of a thiourea linkage. The hydrophobic aromatic fluorescein molecule can be considered as a model small molecule drug that could be exploited as an imaging modality. The conjugation of FITC increased the hydrodynamic size to 127 nm with monomodal size distribution (Figure S5A in SI) and the zeta potential decreased from -51.97±8.35 mV to -40.50±1.08 mV (Figure S5B in SI). The lyophilized FITC conjugated HP-AuNPs (FITC-HP-AuNPs) displayed excellent stability even after freeze-drying and could be easily redispersed in PBS buffer without affecting the size distribution.

### *In-vivo evaluation of FITC-HP-AuNPs* biodistribution in the mouse central nervous system in healthy C57BL/6JRj mice

The brain parenchyma can be accessed by passing through the blood-brain barrier or, in some cases, diffusion into the cerebrospinal fluid produced by the choroid plexus.^[39]^ We screened the brains of female C57BL/6JRj mice intravenously injected with 4 mg/kg (100 µg in 200 µL PBS) FITC-HP-AuNPs 3 h post-injection to determine the actual distribution of the nanoparticles within the diencephalon tissue and choroid plexus. We first verified whether the green signal detected in the brain was derived from the FITC-HP-AuNPs, by imaging with an anti-FITC antibody (Figure 4A). We then compared the green fluorescence intensity levels of the non-injected animals (Figure 4B, n = 3) to those of intravenously injected mice (Figure 4C, n = 3). As illustrated in Figure 4C, mice injected with saline did not exhibit detectable levels of fluorescence in the 488-560 nm wavelength (green) in the brain parenchyma (DAPI, blue, arrowheads) or around the choroid plexus blood vessels (podocalyxin, red, arrows). Interestingly, after 3 h, FITC-HP-AuNPs injected animals exhibited comparable fluorescence intensity levels (Figure 4C) to the HP-FTSC conjugate observed in the brain parenchyma after 1 h (Figure 3A). Increased green fluorescence in the choroid plexus also suggests that FITC-HP-AuNPs can penetrate the cerebrospinal fluid, possibly through the choroid plexus blood vessels or *via* the clearance of the nanoparticles from the brain tissue. We further searched the different regions of the diencephalon for the potential preferential accumulation of the FITC-HP-AuNPs into a given cerebral area. The outer parts of the brain, including the grey matter/cerebral cortex (Figure 4D), exhibited visibly stronger green fluorescence intensity compared to the inner brain regions such as the striatum (Figure 4E). However, nanoparticles were detected in both inner and outer regions of the brain parenchyma, furthermore supporting the evidence of the FITC-HP-AuNPs’ passage through the BBB.

**Figure 4.**
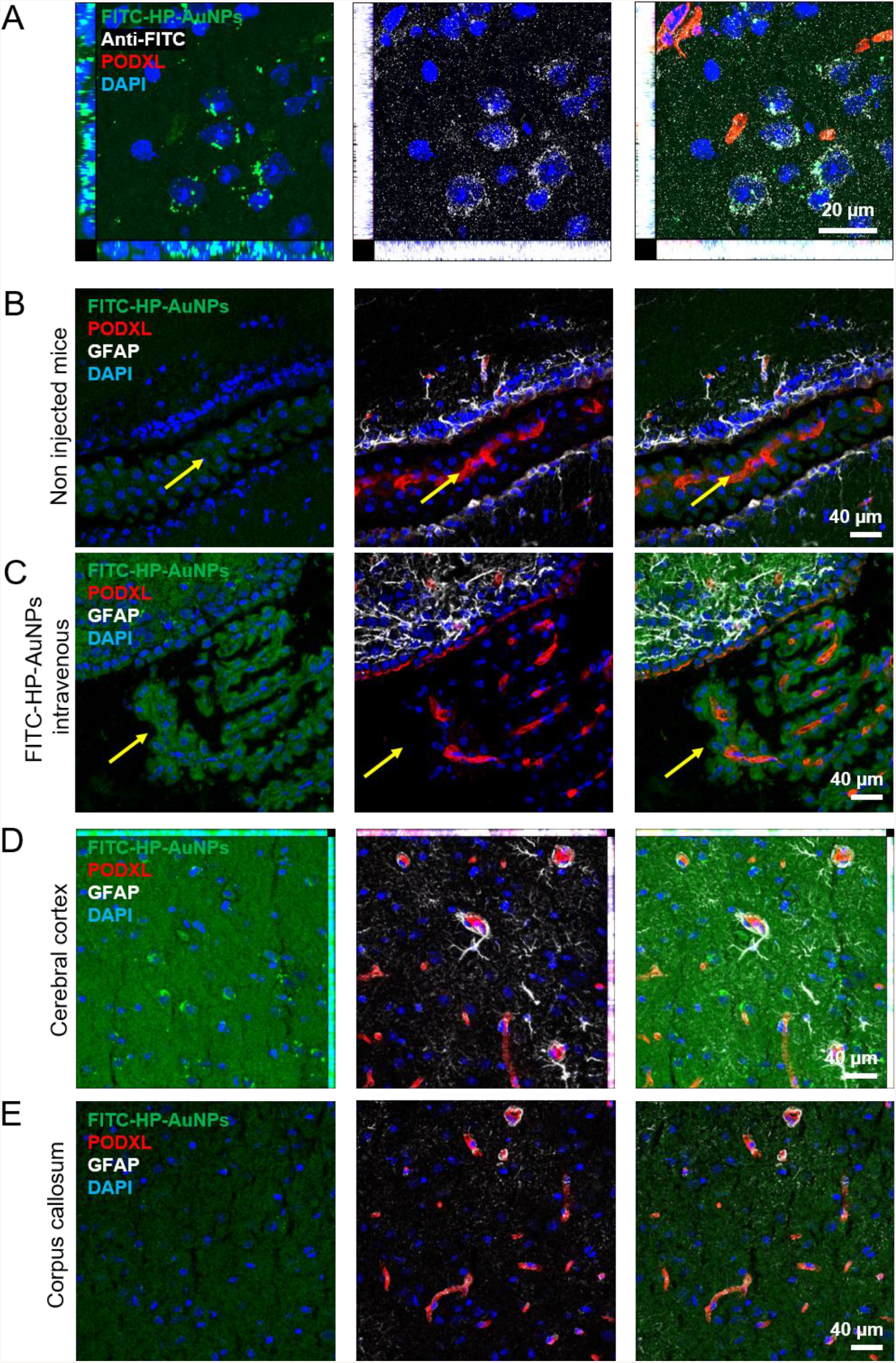
Distribution FITC-HP-AuNPs in the murine central nervous system after peripheral infusion through the caudal vein. **A**, C57BL/6JRj female mice brain micrographs of the immunofluorescent labeling of brain endothelial cells (PODXL, red), FITC-HP-AuNPs (green), and an antibody directed against FITC (white). **B**, Brain micrographs of a C57BL/6JRj female mice injected with saline (PBS, 200 µL) through the caudal vein. After 3 h the brain was collected, immunofluorescent labeling of the endothelial podocalyxin (PODXL, red), astrocytic glial fibrillary acidic protein (GFAP, white), and DAPI nuclear counterstaining (blue) were performed. The yellow arrow indicates the location of the choroid plexus. **C**, Brain micrographs of a C57BL/6JRj female mice injected with 4 mg/kg FITC-HP-AuNPs (100 µg in PBS, 200 µL) through the caudal vein. After 3 h the brain was collected, immunofluorescent labeling of the endothelial cells (PODXL, red), astrocytes (GFAP, white), and DAPI nuclear counterstaining (blue) were performed. The yellow arrow indicates the location of the choroid plexus, the arrowheads highlight intraparenchymal nanoparticles. **D**, Brain micrographs of the cerebral cortex of a C57BL/6JRj female mice injected with 4 mg/kg FITC-HP-AuNPs (100 µg in PBS, 200 µL) in the caudal vein. After 3 h the brain was collected, immunofluorescent labeling of the endothelium (PODXL, red), astrocytes (GFAP, white), and DAPI nuclear counterstaining (blue) were performed. **E**, Brain micrographs of the corpus callosum of a C57BL/6JRj female mice injected with 4 mg/kg FITC-HP-AuNPs (100 µg in PBS, 200 µL) through the caudal vein. After 3 h the brain was collected, immunofluorescent labeling of the endothelial podocalyxin (PODXL, red), astrocytes (GFAP, white), and DAPI nuclear counterstaining (blue) were performed.

### *In-vivo* evaluation of preferential accumulation of HP-AuNP-FITC in different brain cells in three different strains of healthy mice

We then investigated whether the increased detection of FITC-HP-AuNPs fluorescence in the brain gray matter compared to the white matter could be associated with the specific binding of the nanoparticle to one brain cell type. Using specific antibodies, we labeled brain endothelial cells (PODXL, Figure 5A-D), the neuronal cell adhesion molecule (NCAM, Figure 5A), nerves/axons expressing the neuropilin-1 (NRP-1, Figure 5B), astrocytes (GFAP, Figure 5C), and the microglial allograft inflammatory factor-1 (IBA-1, Figure 5D). We performed comprehensive screening of brain biodistribution in three different mice strains, including both male and female individuals, namely immunocompetent female C57BL/6JRj and male FVB mice in addition to the immunocompromised female Rj: NMRI-Foxn1nu/nu (referred to as NMRI-Nude) strain. These mice were intravenously injected with 4 mg/kg FITC-HP-AuNPs (100 µg in 200 µL PBS) and analyzed after 3 h. It emerged that, regardless of the mouse strain, FITC-HP-AuNPs seemed to be preferentially distributed in the brain extracellular matrix with no clear cellular association (Figure 5E). However, the FITC signal from FITC-HP-AuNPs was occasionally found to overlap with the immunofluorescence of identified cells such as neurons or microglial cells, but without any significant association to a particular cell type in any of the three tested mouse strains (Figure 5).

**Figure 5.**
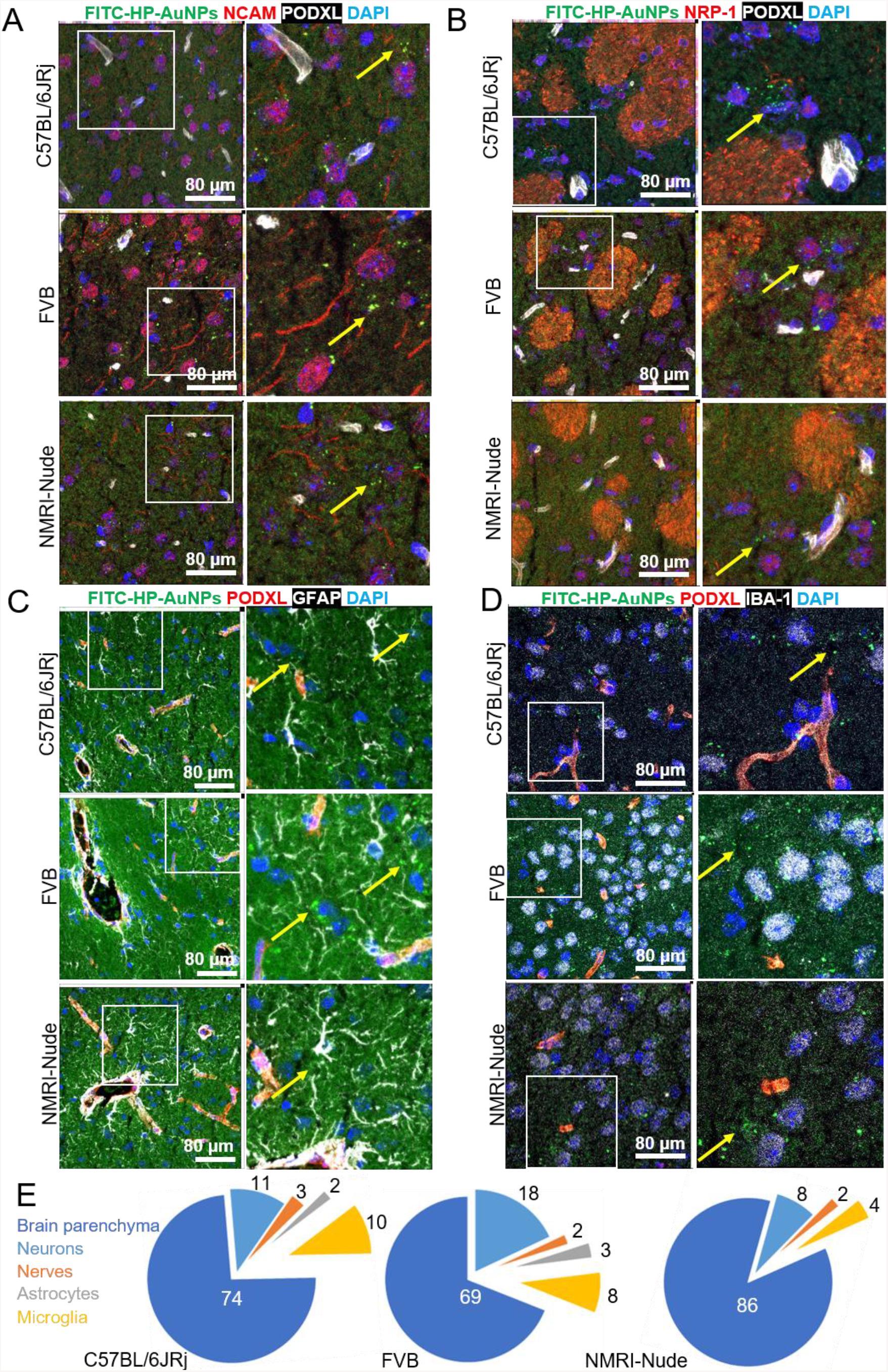
FITC-HP-AuNPs are evenly distributed in the brain tissue and did not home to specific cell types. **A**, Brain micrographs of female C57BL/6JRj, male FVB, and female NMRI-Nude mice injected intravenously with 4 mg/kg of FITC-HP-AuNPs (100 µg) (green) for 3 h. Neurons were labeled using an NCAM antibody (red) and brain endothelial cells with a PODXL antibody (white) 3 h post-injection. Arrows are pointing toward the classical distribution of the nanoparticles, e.g., found isolated in the brain parenchyma. **B**, Brain micrographs of female C57BL/6JRj, male FVB, and female NMRI-Nude mice injected intravenously with 100 µg of FITC-HP-AuNPs (green) for 3 h. Pack of axons/nerves from the striatum were labeled using an NRP-1 antibody (red) and brain endothelial cells with a PODXL antibody (white). Arrows are pointing toward the classical distribution of the nanoparticles, e.g., found isolated in the brain parenchyma. **C**, Brain micrographs of female C57BL/6JRj, male FVB, and female NMRI-Nude mice injected intravenously with 100 µg of FITC-HP-AuNPs (green) for 3 h. Astrocytes were labeled using a GFAP antibody (white) and brain endothelial cells with a PODXL antibody (red). Arrows are pointing toward the classical distribution of the nanoparticles, e.g., found isolated in the brain parenchyma. **D**, Brain micrographs of female C57BL/6JRj, male FVB, and female NMRI-Nude mice injected intravenously with 4 mg/kg (100 µg) of FITC-HP-AuNPs (green) for 3 h. Both peripheral and resident microglia were labeled using an IBA-1 antibody (white) and brain endothelial cells with a PODXL antibody (red). Arrows are pointing toward the classical distribution of the nanoparticles, e.g., found isolated in the brain parenchyma. **E**, Ratios between cell-associated and extracellular/brain parenchyma-associated FITC-HP-AuNPs calculated from the colocalization of fluorescence in the brain micrographs using CellProfiler. Ratios are expressed as the average percentage of all nanoparticles found on ten confocal fields per mouse, with n = 3 for each mouse strain.

### *In-vitro* evaluation of the mechanism of BBB penetration in murine microvascular brain endothelial cells

Intrigued by this result, we intended to study the mechanism of BBB penetration by *in-vitro* experiments using murine microvascular brain endothelial cells (bEND3). We first tested if the incubation of HP-AuNP with bEND3 cells for 24 h alters the expression of gap junction proteins and other BBB transporters, namely, ZO-1, Claudin 5, Glut-1, and Transferrin receptor by real-time quantitative reverse transcription-polymerase chain reaction (RT-qPCR). However, we did not observe any downregulation of the tested protein expressions after 24 h incubation with HP-AuNPs (Figure S9 A in SI), suggesting that the BBB passage of HP-Au-NPs is not due to the disruption of BBB. This is in agreement with the recent study which indicated that HP ameliorates the leaky brain endothelium by restoring the injured glycocalyx in mice.^[40]^ Next, we investigated if the uptake of HP-AuNPs by the bEND3 cells follows an energy-dependent receptor-mediated uptake. For this purpose, we performed the cellular uptake study of FITC-HP-AuNPs at 4 °C by flow cytometry. Interestingly, we observed a ∼80% decrease in uptake of FITC-HP-AuNPs under these conditions suggesting the role of an unknown heparin-sensitive cell-surface receptor (Figure S9 B in SI).

### Synthesis and characterization of radiolabeled ^68^Ga-HP-AuNPs

Next, we intended to quantitatively determine the amounts of HP-AuNPs penetrating the BBB and its biodistribution in different organs. To investigate this, we decided to radiolabel HP-AuNPs with ^68^Ga^3+^ as the radioactive nuclei and examine the kinetics of biodistribution in healthy rats using live positron emission tomography (PET). Larger animals such as rats exhibit a bigger central nervous system than mice which enables studies of the selective binding to specific brain regions with a better resolution. In addition, obtaining data in rats would assist us to evaluate the variability across species which is vital for establishing the versatility of the delivery agent. To develop ^68^Ga complexed HP-AuNPs, we conjugated isothiocyanate functionalized macrocyclic chelator DOTA (1,4,7,10-tetraazacyclododecane-1,4,7,10-tetraacetic acid) using the same protocol as optimized for FITC conjugation. The macrocyclic chelator DOTA is well known to chelate ^68^Ga, analogous to ^111^In-DTPA labeling of heparin conjugate^[41,42]^. As anticipated, the DOTA functionalized HP-AuNPs were obtained with 3% modification (with respect to the carboxylate groups), which was complexed with ^68^Ga^3+^. However, while we could separate a labeled high molecular weight fraction from unbound free ^68^Ga^3+^ using size exclusion chromatography, the majority of the activity was lost after challenging the HP-AuNPs-^68^Ga with the strong chelator EDTA (thermodynamic stability constant log KML = 21.0).^[43]^ This indicates that the majority of the radioactivity associated with the high molecular weight fraction was due to ionically bound ^68^Ga^3+^ with the negatively charged sulfate and carboxylate moieties in the heparin backbone. We, therefore, decided to continue with a chelator–free labeling approach using HP-AuNPs without DOTA, exploiting the excess sulfates and carboxylates on the NP surface. An advantage of this approach is that the radionuclide can be incorporated into the nanoparticle in a single step with no modifications of the HP-AuNPs. However, the risk of radionuclide desorption is a potential drawback and from a chemical point of view transchelation of bound ^68^Ga^3+^ from the HP-AuNP to transferrin (log KML=20.3)^[44]^ may take place in the plasma.

In order to eliminate the risk of false-positive readout by potential desorption of gallium from HP-AuNPs, it is imperative to use a control that could mimic the unbound “free” ^68^Ga^3+^ in the *in-vivo* system. However, while ^68^Ga^3+^ will be stable in its free hydrated gallium ion [Ga(H_2_O)_6_]^3+^ form in the low pH 0.1 M hydrochloric acid generator eluate, it has rather complex chemistry at higher pH and the literature is not entirely consistent. When the pH is increased several neutral or charged hydroxides may be formed and they exist in equilibrium; Ga_6_(OH)_15_, Ga_4_(OH)_11_, neutral insoluble amorphous Ga(OH)_3_, insoluble GaO(OH), Ga_3_(OH)_11_ and soluble gallate Ga(OH)^4-^. The consensus is that the rate of formation of hydroxide species is likely dependent not only on pH but also on concentration, temperature, counterions, and time. At carrier-free conditions precipitation of Ga(OH)_3_ begins at pH ∼3 in the absence of stabilizing ligands. At pH 7.4 the predominant species is insoluble amorphous Ga(OH)_3_. The Ga(OH)_3_ converts on aging to the stable crystalline phase GaO(OH), which is somewhat less soluble in neutral solution than Ga(OH)_3_.^[45]^ Gallium can also form highly insoluble phosphates such as GaPO_4_ at pH values close to neutral hence we avoided the use of any phosphate buffers for the preparation of our “free” gallium control.^[46]^ Instead, we decided to neutralize the hydrochloric acid eluate using an equimolar amount of NaOH and simultaneously formulate our control at pH 4.6 using acetate buffer. The acetate ion is known to be a weak gallium coordinator and can serve to stabilize the free ^68^Ga^3+^ in solution at increased pH and potentially slow down the formation of insoluble ^68^Ga (OH)_3_/^68^GaO(OH). At the same time, we also wanted to minimize the time between synthesis and animal injection, hence we prepared the ^68^Ga-acetate from the hydrochloric acid generator eluate less than five minutes before injection.

### *In-vivo* plasma stability studies of ^68^Ga-HP-AuNPs in Sprague Dawley rats

We injected the freshly prepared ^68^Ga-acetate or ^68^Ga-HP-AuNPs via tail vein in Sprague Dawley rats and performed PET/MRI imaging, quantitative evaluation of plasma stability of NPs, and ex vivo autoradiography binding assay. We first investigated the presence of low molecular weight ^68^Ga-species in the rat plasma samples after administration of ^68^Ga-HP-AuNPs and ^68^Ga-acetate (Figure S12 in SI). Throughout the scan, we observed that around 90% of the plasma radioactivity was associated with the high molecular weight (HMW) fraction (>5 kDa) after the injection of ^68^Ga-HP-AuNPs. The ^68^Ga-acetate, on the other hand, was rapidly associated with plasma proteins (approximately 70%), presumably via transferrin binding. We could detect a small amount of low molecular ^68^Ga-species in the plasma in the ^68^Ga-HP-AuNP case (<10%). Similar to the weakly chelated ^68^Ga-citrate, we speculate that only the soluble gallate ^68^Ga(OH)^4−^ is formed *in-vivo* because the HP-AuNP can prevent precipitation of ^68^Ga(OH)_3_. Any desorbed ^68^Ga^3+^ from the HP-AuNP is likely to immediately form the small charged molecule ^68^Ga(OH)^4−^ which may exhibit rapid renal elimination. However, ^68^Ga(OH)^4−^ may also potentially be taken up by transferrin. ^[47]^ On the other hand, ^68^Ga-Transferrin and ^68^Ga(OH)^4−^ exist in equilibrium in plasma, and unbound ^68^Ga(OH)^4−^ will continuously be excreted through urine, which presumably explains the increased radioactive accumulation in the bladder.

PET imaging demonstrated rapid blood clearance of the ^68^Ga labeled HP-AuNPs from the circulation over the first 90 minutes after administration (estimated from segmentation of the left heart ventricle, or by measurement of sampled blood and plasma), consistent with the size of the construct. Bladder accumulation was observed for ^68^Ga-HP-AuNP (Table 2) but not ^68^Ga-acetate, confirming that ^68^Ga^3+^ desorbed from ^68^Ga -HP-AuNP, in fact, was excreted through urine, presumably in the form of ^68^Ga(OH)^4−^ as postulated above, and not bound to transferrin. Thus, the limited urinary excretion and plasma data together confirmed the stability of the ^68^Ga labeled HP-AuNPs.

**Table 2.**
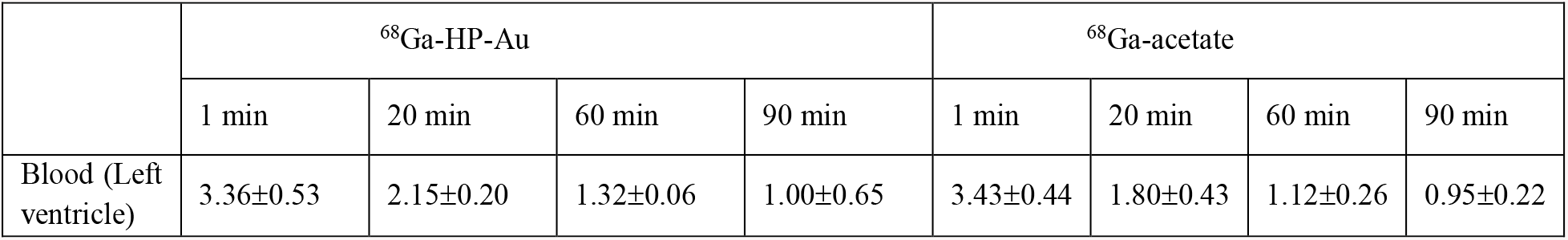
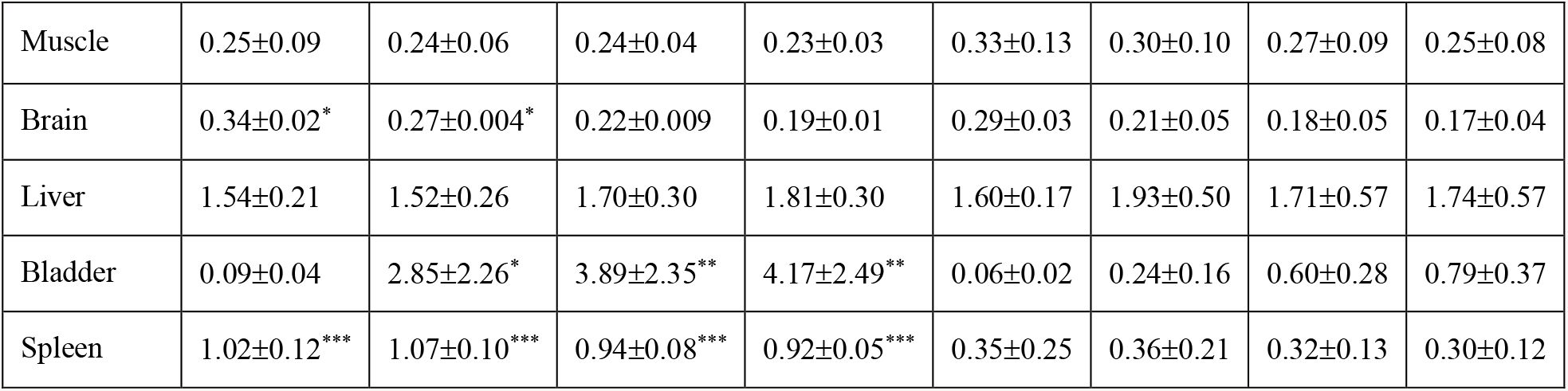
The difference in biodistribution between ^68^Ga-HP-Au the CNS and 68Ga-acetate, where tissue uptake is expressed as %ID/g. Statistical differences (unpaired t-test) indicated by stars, where * indicates p<0.05, ** indicates p<0.01, and *** indicates p<0.001.

### *In-vivo* PET/MRI evaluation and biodistribution study of ^68^Ga-HP-AuNPs in Sprague Dawley rats

Next, we focused on the brain distribution of ^68^Ga-HP-AuNP and the non-selective control ^68^Ga-acetate following the tail-vein injection. The biodistribution of ^68^Ga-acetate was different from that of ^68^Ga-HP-AuNPs, especially in the bladder, spleen, and brain (Table 2). Dynamic whole-body imaging (Supplemental Figure S11 and Figure 6A) illustrates the rapid excretion through the bladder (detected from 4 min on) and metabolization by the liver. Image analysis of the brain indicated that the ^68^Ga-HP-AuNP showed faster brain accumulation compared to the ^68^Ga-Acetate especially at earlier time points (0 – 40 min) (Figure 6B). Live imaging indicates a clear and strong signal overlapping with the base arteries irrigating the brain, also named Circle of Willis (Figures 6C-E).^[48]^ The accumulation of ^68^Ga-HP-AuNP in the brain was approximately 0.30% ID/g shortly after administration and 0.20 %ID/g after 90 minutes, which was, as a reference, around half the accumulation to the splenic tissue (Table 2).

**Figure 6.**
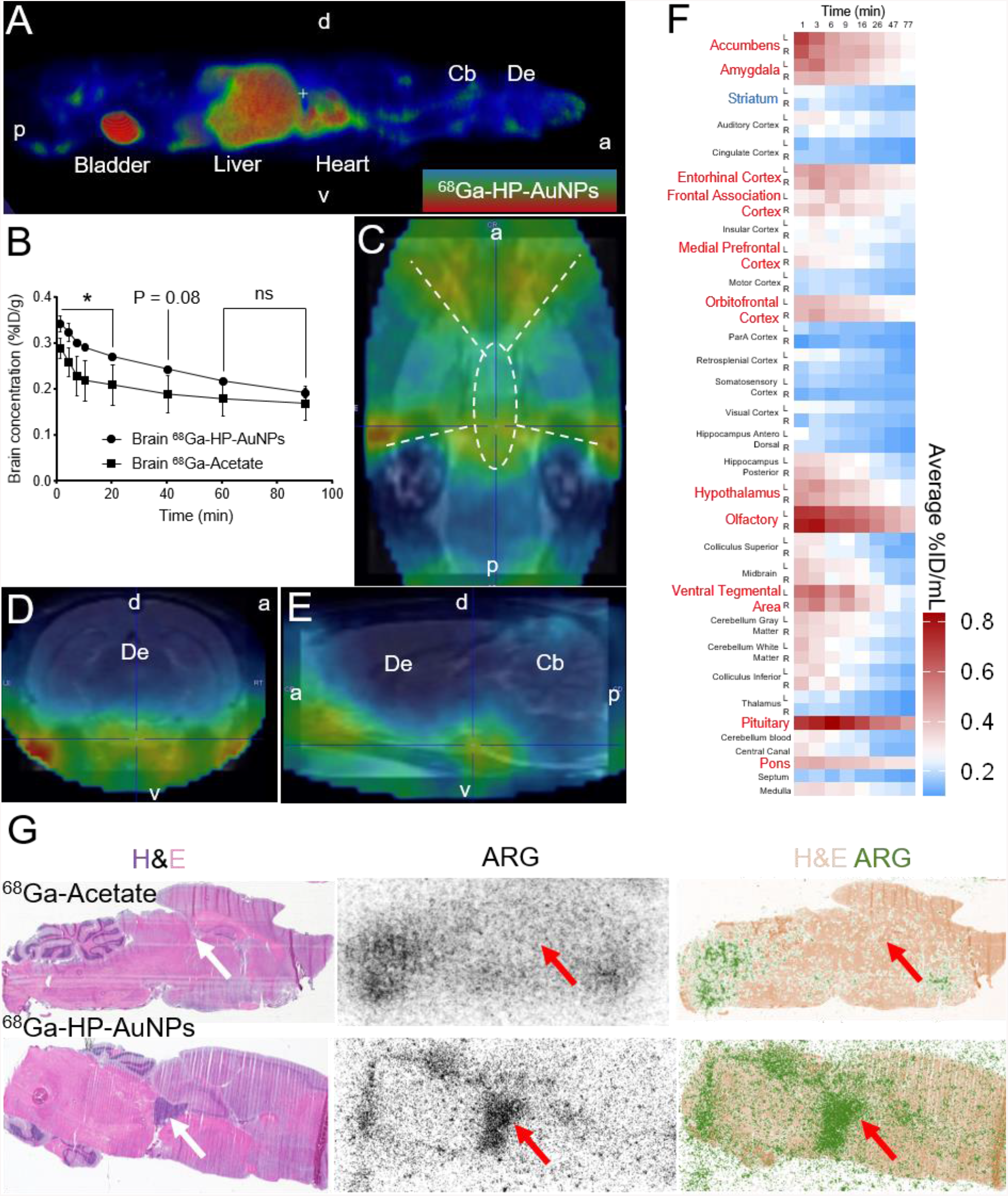
Rat central nervous system distribution of ^68^Ga-HP-AuNP. **A**, Representative whole-body dynamic PET images seven minutes after administration of ^68^Ga-HP-AuNP, normalized to standard uptake value (SUV) 10. p = posterior, a = anterior, d = dorsal, v = ventral, Cb = cerebellum, De = diencephalon (right). **B**, Increased delivery of ^68^Ga-HP-AuNP (circle points) to the CNS compared to ^68^Ga-acetate (square points). n = 2. *, p < 0.05, n.s = non-significant. **C-E**, Coronal ventral section of the brain (**C**), the trans-axial section of the diencephalon (**D**), and sagittal lateral section (**E**) point out increased radioactivity in the Circle of Willis (dotted lines on (C)), i.e., the base arteries of the central nervous system after the administration of ^68^Ga-HP-AuNP (summation image of all time points 0-90 minutes post-administration). p = posterior, a = anterior, d = dorsal, v = ventral, Cb = cerebellum, De = diencephalon. **F**, Heatmap of the supervised PMOD-FUSEIT T2-weighted MRI image analysis aligned on the Rat brain Atlas coordinates for the quantitation of ^68^Ga-HP-AuNP in the left (L) and right (R) hemispheres at the indicated time points of the live imaging. n = 2. **G**, Rat brain histological section counterstained with hematoxylin & eosin (H&E, left), autoradiography of the same section (middle), and merged pictures (right) for rats injected with ^68^Ga-acetate (top) or ^68^Ga-HP-AuNP (bottom). Unlike the ^68^Ga-acetate, increased ^68^Ga-HP-AuNP radioactivity is observed in the ventricular spaces (arrows). n = 3.

The total uptake of ^68^Ga-HP-AuNPs in the brain was 0.25 %ID, which translates to exposure of 1.25 µg of HP-AuNPs (since 500 µg was administered in total). This difference between the ^68^Ga-HP-AuNP and the ^68^Ga-acetate tends to disappear beyond 40 min, suggesting a slow and passive diffusion of the ^68^Ga-acetate conjugate as opposed to the potential active transportation and/or specific binding to a BBB transporter of the ^68^Ga-HP-AuNPs, allowing its increased diffusion rate to the brain (Figure 6B). To compare the mouse brain fluorescence distribution data obtained with the FITC-HP-AuNPs (Figure 4 and 5), we performed an extensive study of the ^68^Ga-HP-AuNPs distribution in the rat brain areas over the course of the live imaging. We used the PMOD-FUSEIT T2-weighted MRI image analysis aligned on the Rat Brain Atlas to determine the amount of ^68^Ga-HP-AuNPs in the left (L) and right (R) brain structures and summarized the data as a heatmap (Figure 6F). In accordance with our observations in mice (Figure 4), the striatum showed lower accumulation of the ^68^Ga-HP-AuNP-derived radioactivity compared to outer cortical structures such as the entorhinal-, frontal associative-, prefrontal- and orbitofrontal cortex (Figure 6F). Interestingly, brain areas such as the pituitary, pons, and olfactory bulbs showed higher and prolonged ^68^Ga-HP-AuNP accumulation (Figure 6F). We speculate that increased blood flow in these regions^[49]^ due to maintained neuronal activity even during anesthesia^[50,51]^ could explain the readout. In accordance with the *in vivo* imaging, *ex vivo* autoradiography imaging of the rat brain sections indicated the presence of the ^68^Ga-HP-AuNP inside the brain parenchyma and in areas containing cerebrospinal fluid, such as the lateral ventricles near the hippocampus, the cerebellum, and brain stem (Figure 6G). The ^68^Ga-acetate did not show a clear association with ventricule-associated structures (Figure 6G). Although the sensitivity and spatial resolution of the autoradiography technique do not match the confocal fluorescence imaging data previously obtained in mice (Figure 5), both methodologies and animal models indicate that the HP-AuNPs can very rapidly and widely penetrate the central nervous system parenchyma and cerebrospinal fluid compartments. This finding offers untapped potential as cargo for the delivery of imaging probes and therapeutic molecules.

### Synthesis and characterization of gadolinium-labeled gold nanoparticles (Gd-HP-AuNPs)

Encouraged by the efficient complexation and delivery of radiolabeled ^68^Ga-HP-AuNP, we envisioned developing MRI-active gadolinium labeled HP-AuNPs. Unlike ^68^Ga^3+^ complexation, which requires acidic pH, the Gd^3+^, could be complexed under basic conditions, which could promote efficient complexation to the sulfated NPs. Upon complexation, the hydrodynamic size of the HP-AuNPs (Gd-HP-AuNPs) reduced from 86 nm to 75 nm (Figure S3 and S7 in SI), while the zeta potential, which represents the surface charge on the HP-AuNPs reduced from - 51.97±8.35 mV to -32.37±6.27 mV (Figure S6B in SI). The particles were highly stable after complexation and did not show any signs of aggregation in saline or PBS buffer. We measured the concentration of Gd and Au in Gd-HP-AuNPs by inductively coupled plasma mass spectrometry (ICP-MS). The ICP-MS study revealed that the concentration of complexed Gd and Au in the Gd-HP-AuNPs were 8 weight% and 3.5 weight% respectively. In order to ascertain the efficiency of the complexation of Gd^3+^ to the sulfates and carboxylates on the HP-AuNP surface, we performed an Arsenazo-III assay.^[52]^ Corroborating with our ^68^Ga^3+^ complexation data, this study revealed that only 1.7% uncomplexed “free” Gd^3+^ was present in Gd-HP-AuNPs (Figure S8B in SI), which suggest strong complexation with the free sulfates and carboxylates, thus, avoiding the need for conventional macrocycles like DOTA. To further ascertain the Gd complexation, we characterized them using FTIR spectroscopy (Figure S5 in SI). All four samples (HP, HP-Dopa-DTPH, HP-AuNPs, and Gd-HP-AuNPs) showed a broad peak at 3100-3600 cm-1 owing to the intramolecular and intermolecular stretching vibrations of hydroxyl groups. The sp3 C-H stretching at 2953 cm-1, the symmetrical stretching vibrations of the S=O group in the NH-SO3-at 1421 cm-1, the asymmetrical stretching vibrations of the S=O group in the CH2-SO3-at 1216 cm-1 and the stretching of -C=O of the carboxyl group at 1614 cm-1 typically signifies HP structure.[53] The C=O stretching peak at 1614 cm-1 shifted to 1640 cm-1 after conjugation of dopamine moiety indicating covalent grafting of dopamine by amide bond. Two new peaks, secondary amide NH bending at 1525 cm-1 and aromatic OH stretching of catechol moiety at 1370 cm-1 further confirm the successful conjugation.[54] During the gold nanoparticle formation, we have observed a shift of aromatic OH stretching of catechol moiety from 1370 cm-1 to 1378 cm-1, suggesting their role in reducing gold and stabilizing the nanoparticles. Otherwise, the secondary amide NH bending shifted to 1500 cm-1 and the C=O stretching peak also shifted to 1653 cm-1. During the gadolinium complexations on the HP-AuNPS, the C=O stretching peak of the carboxylates significantly shifted to 1632 cm-1. The sulfate peaks, i.e. the symmetrical stretching vibrations of the S=O group in the NH-SO3- and the asymmetrical stretching vibrations of the S=O group in the CH2-SO3-also shifted to 1429 and 1210 cm-1 respectively, suggesting the complexation of Gd ion via the carboxylates and sulfates of the polymer backbone. We also performed the thermal analysis of the samples to confirm the amount of Gd complexation into the samples (Figure S8A in SI). The typical mass loss (∼15%) was observed between the 40 °-200 ºC range attributed to the loss of physisorbed water molecules. The steep decline in mass within the range of 200 °C - 650 ºC is attributed to the oxidation of the polymer backbone along with the loss of the functional groups. The effective remaining residues at 950 °C indicated the presence of 8% Au and 9.8% Gd in Gd-HP-AuNPs. The %Gd result obtained by TGA analysis is in close agreement to that obtained from ICP-MS analysis. We believe, the higher residual mass after TGA analysis is due to the affinity of HP polymer to complex with metal salts.

### *In-vitro* relaxivity study

We first evaluated the relaxation properties of our formulated nanocarrier used as T1-weighted contrast agents in an *in-vitro* MRI study. T1-weighted spin-echo MRI at 1T also revealed the contrast enhancement of our formulated nanocarrier compared to the equivalent concentrations of commercial contrast agent, Gd-DTPA (Figure 7A). The relaxivity of Gd-HP-AuNP was 9.89 mM^- 1^s^-1^, which is ∼3 fold higher than that of commercial Gd-DTPA (3.26 mM^-1^s^-1^) (Figure 7B).^[55]^ The increased longitudinal relaxation of trapped Gd^3+^ ions resulted in contrast enhancement. The presence of heparin macromolecule would presumably have a major influence on the inner and outer coordination sphere hydration, resulting in slow rotational diffusion of the MRI agent.^[56]^

**Figure 7.**
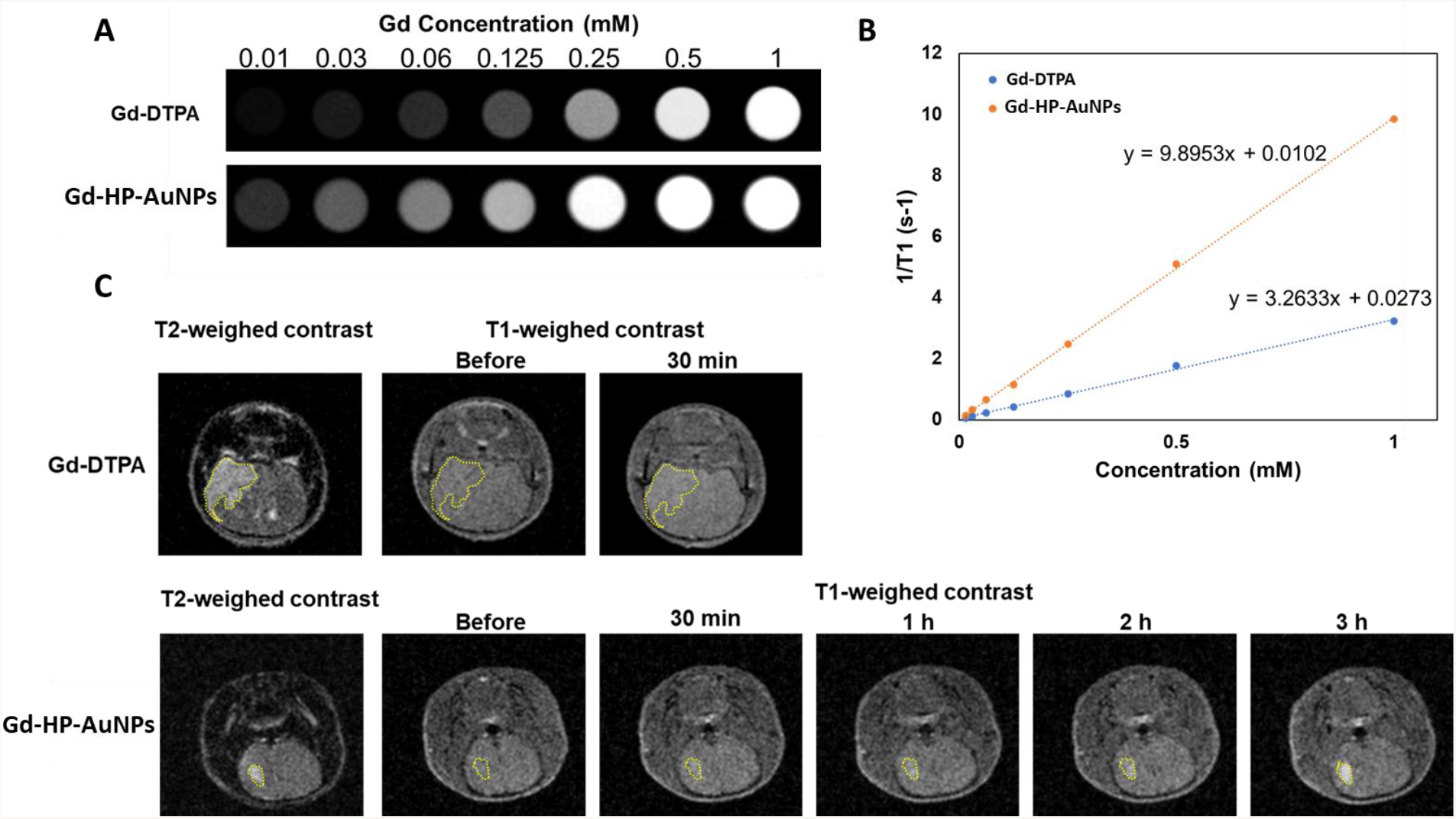
MRI study of Gd-HP-AuNPs. **(A)** T1-weighed images of commercial contrast agent and HP-Au-Gd nanoparticles at different Gd concentrations. **(B)** The T1 relaxation rate of Gd-DTPA and Gd-HP-AuNPs at different Gd concentrations. **(C)** *In-vivo* MR images of intracranial U87-MG tumor-bearing mice before and after intravenous injection of Gd-DTPA (upper panel) and Gd-HP-AuNPs (lower panel). T2-weighed images were used to locate the tumors. T1-weighed imaging was used to detect the contrast enhancement.

### *In-vivo* evaluation of Gd-HP-AuNP localization in a glioblastoma model

Finally, we injected the Gd-HP-AuNPs through the tail vein in nude mice (female BALB/c, 6 weeks old) bearing orthotopically implanted U87MG tumors, and the MRI reading was recorded. The T2-weighed images were initially used to locate the tumors before injecting the MRI probes. Then, the contrast enhancement of intracranial T1-weighed images was used to trace the accumulation of the clinical agent Gd-DTPA and Gd-HP-AuNPs in the tumors. Gd-DTPA showed slight contrast enhancement in the tumor at 30 minutes after injection (Figure 7C). Because Gd-DTPA is rapidly cleared from the body, we did not image longer times. ^[57]^ Gd-HP-AuNPs showed a gradual increase in the contrast in the tumor region between 1 h to 3 h (Figure 7C), indicating successful accumulation in the xenografts. Though the exact mechanism of NP accumulation to the glioblastoma in the xenograft model is not fully understood, we believe the heparin NPs target hepatocyte growth factor/scatter factor (HGF/SF) or heparan sulfate proteoglycans that are overexpressed on GBM.^[58]^ We are currently investigating this further for drug delivery applications which will be communicated elsewhere.

In conclusion, we have discovered a novel biomimetic drug delivery system that can penetrate BBB and distribute it throughout the brain in healthy rodent models but can also target glioblastoma in a xenograft model. The tailored HP-AuNPs could be easily modified with radiotracers (^68^Ga^3+^) and MRI contrast agents (Gd^3+^), without the need for chelating molecules, exploiting the electrostatic interactions between the anionic functional groups (sulfates and carboxylates) in the polymer backbone and the cationic charge on the imaging agents (^68^Ga^3+^, Gd^3+^). The metal complexation on the HP-AuNPs was fast, efficient, and was stable in blood plasma. The *in-vivo* trafficking of the HP-AuNPs to the brain was validated in different immunocompetent (Sprague Dawley rats, C57BL/6JRj, and FVB) as well as in immunocompromised (NMRI-nude) mice models using confocal microscopy, PET, and MRI experiments. The radio tracing experiments in live animals revealed that approximately 0.30%ID/g of the ^68^Ga-HP-AuNPs is accumulated in the brain was shortly after administration and 0.20 %ID/g remained after 90 minutes. These NPs were mainly distributed in the brain parenchyma and areas containing cerebrospinal fluid, such as the lateral ventricles near the hippocampus, around the cerebellum, and brain stem. These results also corroborated with the *ex-vivo* confocal analysis of the brain tissue samples which indicated that the fluorescently labeled FITC-HP-AuNP were uniformly distributed in the brain parenchyma (69-86%) with some accumulation in neurons (8-18%) and microglia (4-10%). The complexation of Gd^3+^ to HP-AuNPs displayed a relaxivity of 9.89 mM^-1^s^-1^, which was ∼3 fold higher than that of commercial Gd-DTPA. The tail-vein administration of Gd-HP-AuNPs in an orthotopic U87MG tumor model showed enriched T1-contrast at the intracranial tumor similar to clinical agent Gd-DTPA. The Gd-HP-AuNPs showed a gradual increase in the contrast in the tumor region between 1 h to 3 h indicating successful accumulation/targeting into brain tumors. The uptake mechanism of HP-AuNPs across the BBB remains to be identified, however, it follows an energy-dependent mechanism without altering the expression of gap-junction proteins as confirmed by the *in-vitro* experiments. We believe, our finding offers HP-NPs as an untapped delivery agent to deliver cargo molecules and imaging probes across the BBB that could be potentially used for diagnostic and therapeutic application, such as for treating neurological disorders.

## AUTHOR INFORMATION

### Author Contributions

The manuscript was written through the contributions of all authors. All authors have approved the final version of the manuscript. ‡These authors contributed equally.

### Notes

Any additional relevant notes should be placed here.

## ACKNOWLEDGMENT

We acknowledge Dr. Ganesh Nawle for his excellent support and assistance in performing the radiolabelling experiment.

