## Supporting Information for "Heparin-derived theranostic nanoprobes overcome the blood brain barrier and target glioma in murine model"

### Methods and Materials

Heparin (HP) from porcine intestinal mucosa (15 kDa), Chondroitin sulfate-A (CS) sodium salt from the bovine trachea (54 kDa), Fluorescein-5-thiosemicarbazide (FTSC), N-hydroxybenzotriazole (HOBt), Fluorescein isothiocyanate isomer I (FITC), N-hydroxysuccinimide (NHS), Dopamine (2-(3,4-Dihydroxyphenyl) ethylamine hydrochloride), 1-ethyl-3-(3-dimenthylaminopropyl) carbodiimide hydrochloride (EDC.HCl), Gold(III) chloride trihydrate ($HAuCl_4$. $3H_2O$) and all other reagents and solvents were purchased from Sigma-Aldrich and used as received. All spectrophotometric analysis was carried out on Shimadzu UV-3600 plus UV-VIS-NIR spectrophotometer.

#### 1. Synthesis and characterizations of nanoparticle systems

##### *1.1 Synthesis of fluorescein thiosemicarbazide (FTSC) conjugated HP and CS nanoparticles*

Fluorescein-5-thiosemicarbazide (FTSC) was conjugated to HP and CS by carbodiimide coupling chemistry using HOBt and EDC following our reported protocol.[1] The orange fluffy materials were obtained after lyophilization and the percentage conjugation was estimated to be 0.5 mol% (with respect to the disaccharide repeat units) by UV-Vis spectroscopy using the

FTSC extinction coefficient of 78,000 $M^{-1}cm^{-1}$ at 492 nm. The $^{1}$H-NMR was not very informative as the conjugation of hydrophobic molecules like FTSC to the HP and CS polymer induces amphiphilicity that promotes self-assembly resulting in the disappearance or underestimation of hydrophobic molecules due to core-shell assembly. The hydrodynamic particle size distribution of HP-AuNPs was carried out using a Zetasizer Nano ZS (Malvern, UK) using 10 × 10 × 45 mm disposable polystyrene cuvette. Freeze-dried samples were dissolved in deionized water at 0.1 mg/mL concentration and stirred at room temperature for 15 minutes before performing the DLS measurement at 25 °C

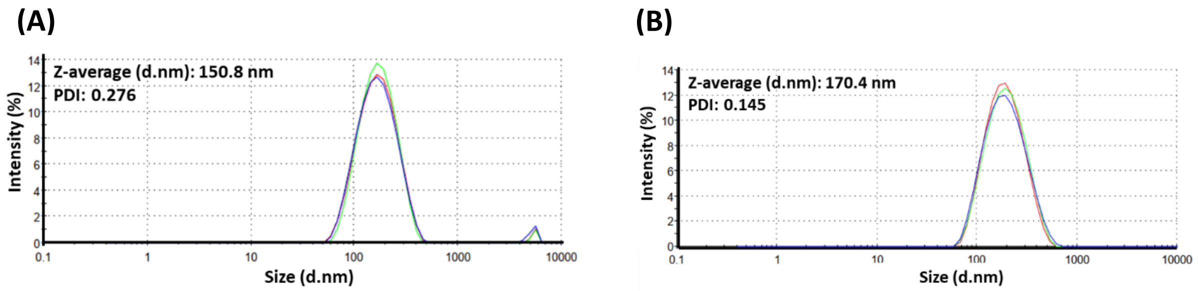


**Figure S1**: Particle size distribution of (A) HP-FTSC and (B) CS-FTSC nanoparticles in water (0.1 mg ml$^{-1}$) as determined by dynamic light scattering measurements.

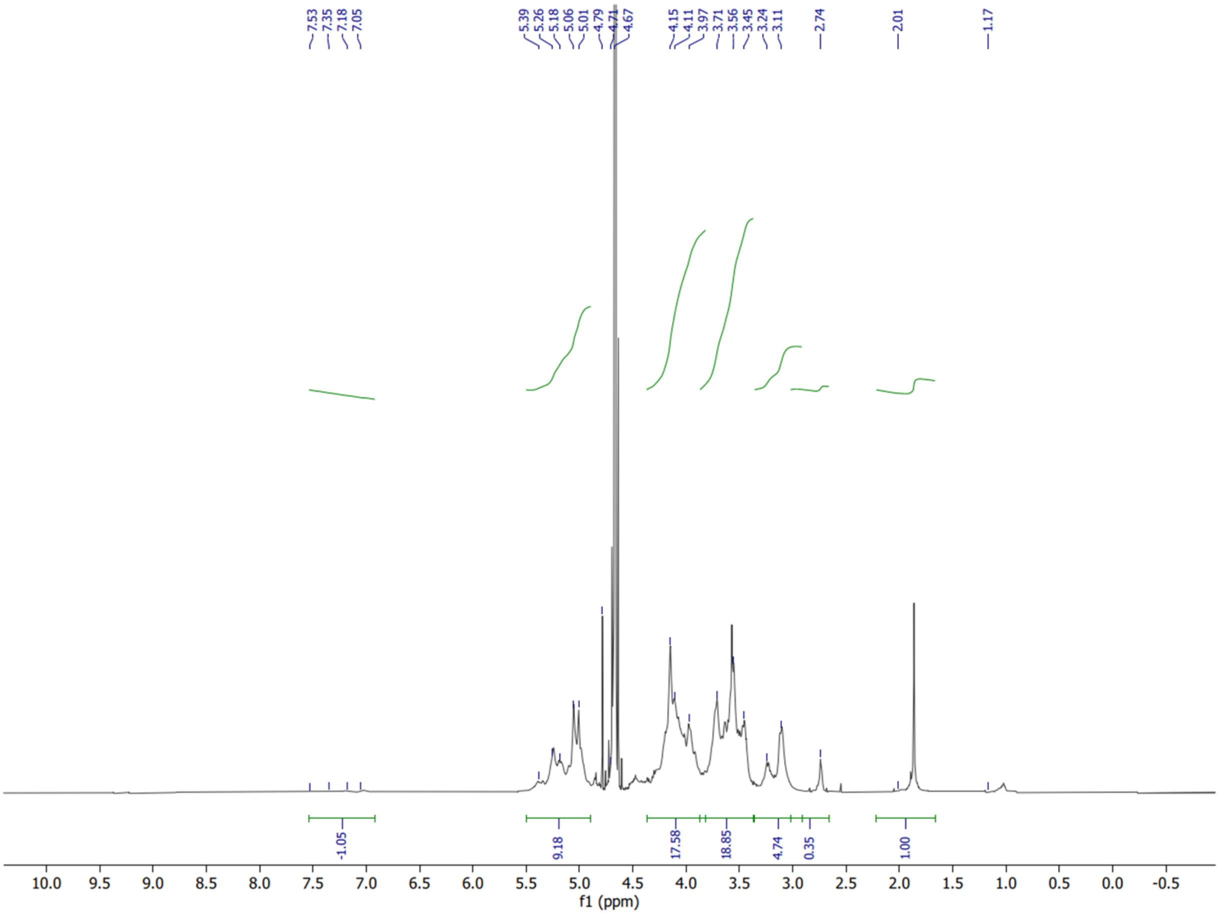

**Figure S1(C)**: [1]H-NMR (500 MHz) spectra of HP-FTSC recorded in $D_2O$ at 298K.

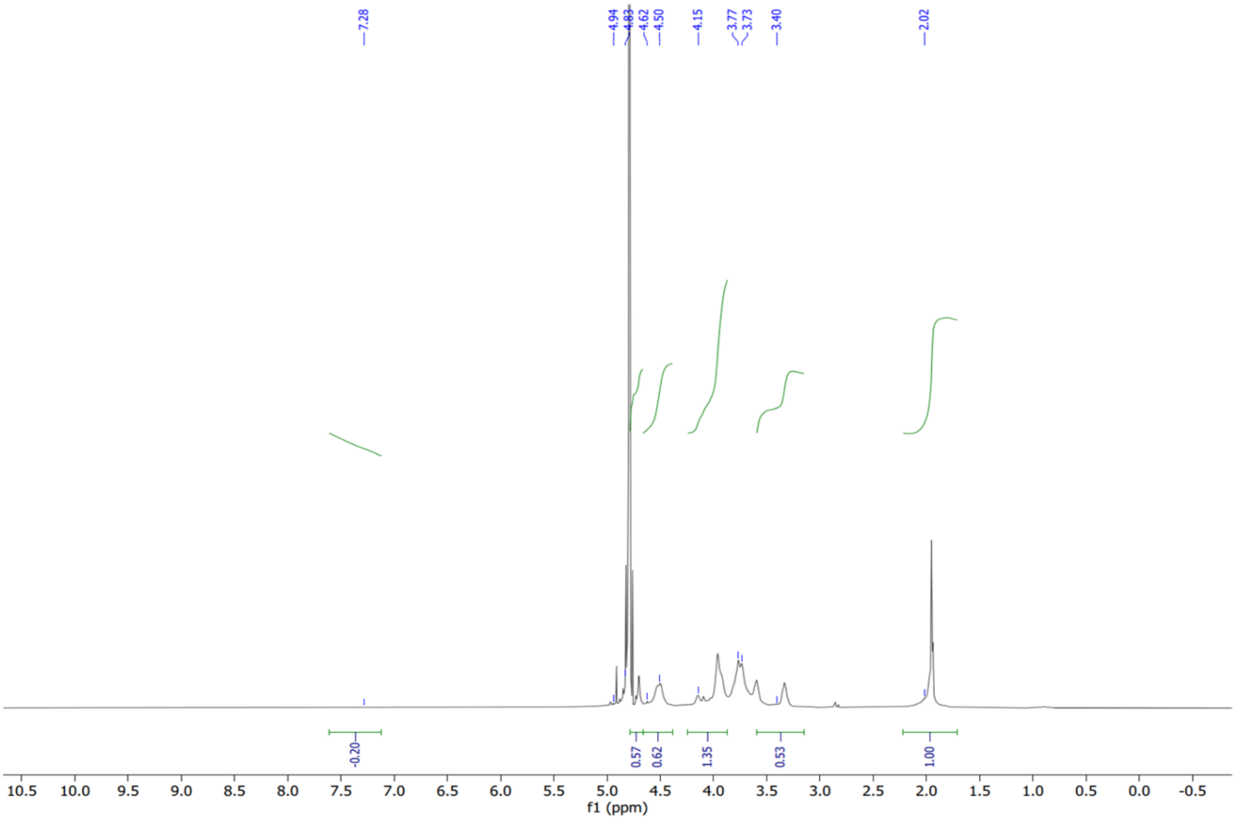


**Figure S1(D):** [1]H-NMR (500 MHz) spectra of CS-FTSC recorded in $D_2O$ at 298K.

#### *1.2 Synthesis and characterization of HP coated gold nanoparticles (HP-AuNPs)*

**Step 1:** Synthesis of HP-Dopa-DTPH

We have conjugated 3,3'-dithiobis (propanoic hydrazide) (DTPH) and dopamine (DA) linker to the carboxylate residues of heparin (HP) in two steps following carbodiimide chemistry to obtain HP-Dopa-DTPH.

Briefly, 0.5 mmol of HP (with respect to disaccharide units) was dissolved in 50 ml of deoxygenated de-ionized water followed by 0.5 mmol of N-hydroxysuccinimide (NHS) and 0.5 mmol of dopamine. Thereafter, the reaction mixture was stirred for 30 min (till it becomes homogeneous) and the pH of the reaction mixture was adjusted to 5.5 by careful addition of 0.1M NaOH. Finally, 0.1 mmol EDC.HCl was added and stirred overnight. The reaction mixture was dialyzed using (Spectra Por-6, MWCO 3500 g/mol) against dilute HCl (pH = 3.5) containing 0.1 M NaCl (4×2L, 48 h), and then against deionized water (3×2L, 24 h).

In the subsequent step, the reaction mixture was transferred back to a 250 mL round-bottom flask. 0.5 mmol HOBt was allowed to dissolve followed by 0.5 mmol of DTPH. The pH of the reaction mixture was adjusted to 4.8 by careful addition of 0.1M NaOH. Finally, 0.25 mmol EDC.HCl was added and stirred overnight. The reaction mixture was dialyzed using (Spectra Por-6, MWCO 3500 g/mol) against dilute HCl (pH = 3.5) containing 0.1 M NaCl (6×2L, 48 h), and then against deionized water (4×2L, 48 h). The solution was lyophilized to obtain fluffy white HA-Dopa-DTPH. The degree of dopamine conjugation was 13 mol% (with respect to the disaccharide units of HA), as estimated by UV-Vis spectroscopy (at pH 7.4 in PBS buffer) using the dopamine extinction coefficient of 2.673 mM$^{-1}$cm$^{-1}$ at 280 nm. From $^{1}$H NMR analysis, the aromatic signal (6.7-6.9 ppm) was 7.7% with respect to the N-acetal signal from the glucosamine units. However, this degree of modification (7.7%) is inaccurate as unfractionated heparin does not possess well-defined repeat units. The degree of hydrazide (DTPH) modifications was determined using trinitrobenzene sulfonic acid (TNBS) assay, which gives 13 mol% modification (with respect to disaccharide repeat units of HA) using UV-Vis spectroscopy. The conjugation of dopamine moiety was further confirmed by the presence of aromatic peaks (6.7-6.8 ppm) in $^{1}$H-NMR (500 MHz) spectra.

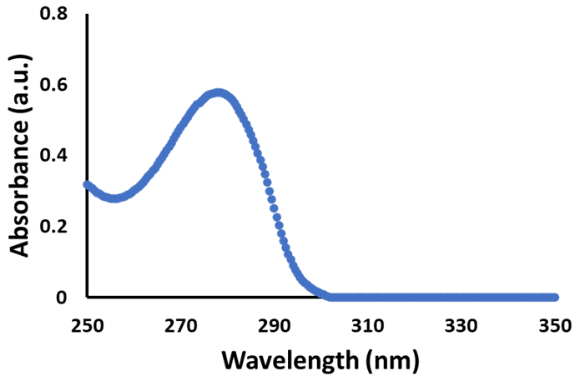


**Figure S2(A):** UV-Vis spectra of conjugated dopamine in HP-Dopa-DTPH dissolved in PBS at 1 mg/mL

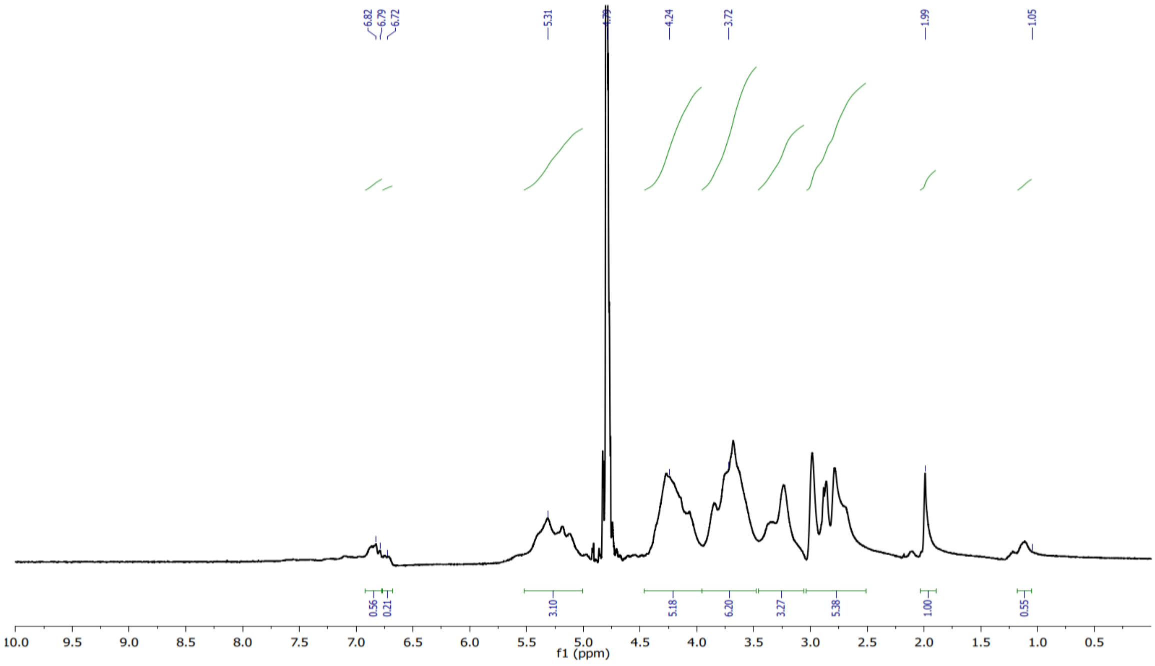

**Figure S2(B):** $^{1}$H-NMR (500 MHz) spectra of conjugated dopamine in HP-Dopa-DTPH recorded in $D_2O$ at 298K.

**Step 2:** Synthesis of HP-AuNPs

HP-Au NPs were synthesized at 40 °C using HP-Dopa-DTPH as both capping and reducing agents following our reported protocol.[2] The fluffy purple-violet material was obtained with the hydrodynamic size of 86 nm (Figure S3) as confirmed by dynamic light scattering (DLS) measurement. The diminishing aromatic peaks of dopamine moiety in the $^{1}$H-NMR spectra indicate the utilization of dopamine in reducing gold chloride to AuNPs.

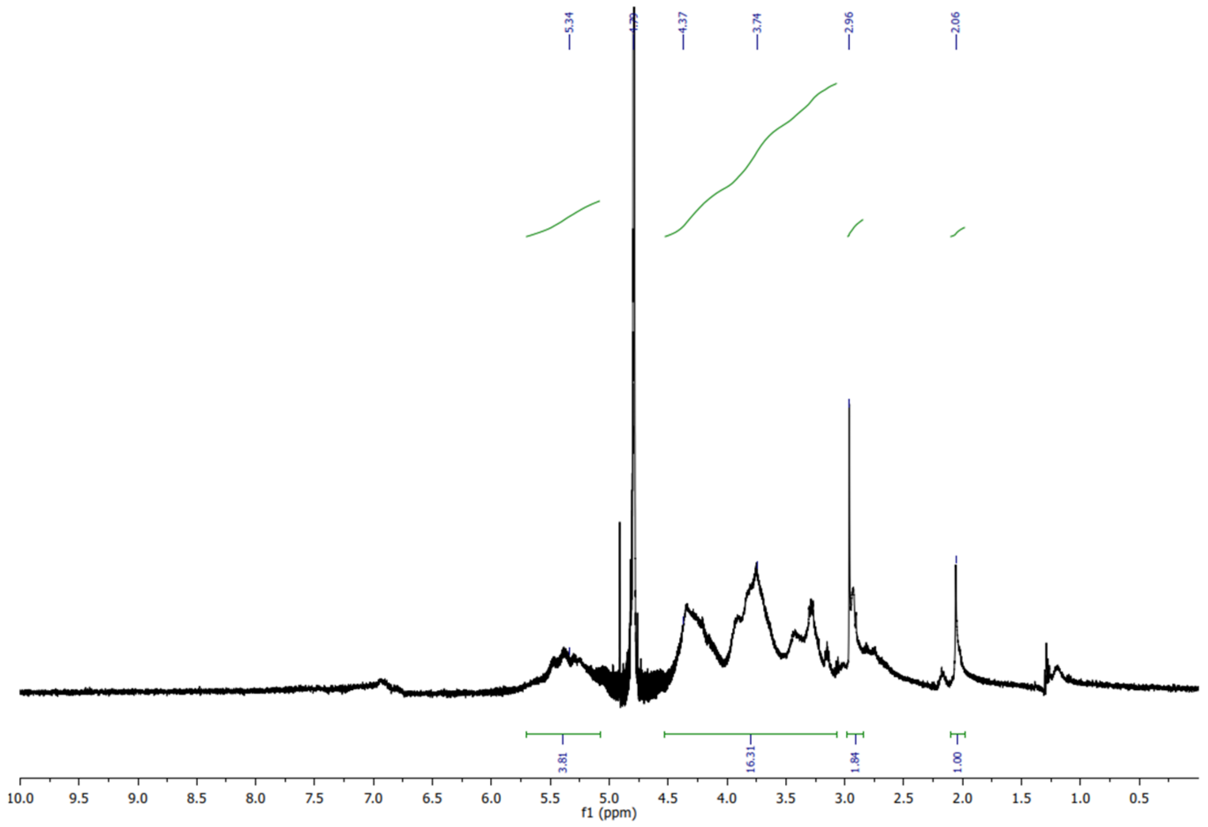


**Figure S2(C)**: $^{1}$H-NMR (500 MHz) spectra of HP-AuNPs recorded in $D_2O$ at 298K.

#### 1.2.1 Hydrodynamic size and surface zeta potential measurement of HP-AuNPs

The hydrodynamic particle size distribution of HP-AuNPs was determined to be 86.67 nm with PDI 0.233 using a Zetasizer Nano ZS (Malvern, UK) using 10 × 10 × 45 mm disposable polystyrene cuvette. Freeze-dried samples were dissolved in deionized water at 0.1 mg/mL concentration and stirred at room temperature for 15 minutes before performing the DLS measurement at 25 °C. The surface zeta potential was subsequently measured to be -51.97±8.35 mV using Zetasizer Nano ZS at 25 °C using disposable folded capillary DTS1070 cells obtained from Malvern, UK.

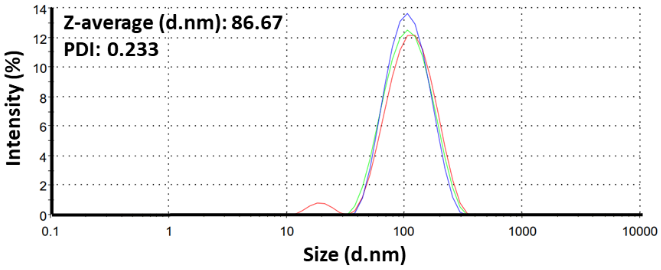


**Figure S3:** Particle size distribution of HP-AuNPs in water (0.1 mg ml$^{-1}$) as determined by dynamic light scattering measurements.

#### 1.2.2 Transmission electron microscopy (TEM)

The morphology of HP-AuNPs was observed by transmission electron microscopy (TEM). The samples were stained with uranyl acetate (10 µL of a 2% w/v solution) for 30 s and set on carbon-coated 400 mesh Cu grids (Nisshin EM). The grids were dried and transferred to an H7000 TEM (Hitachi Ltd, Tokyo, Japan) for imaging at an acceleration voltage of 75 kV.

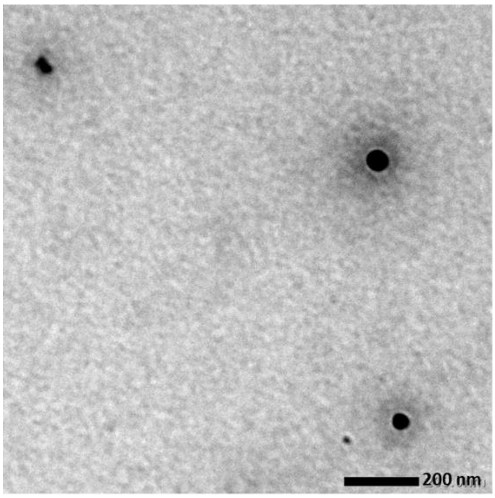


**Figure S4:** Particle size of HP-AuNPs obtained from TEM measurement (scale bar: 200 nm)

#### 1.2.3 Attenuated total reflectance (ATR) FTIR spectroscopy

We have further supported the conjugation of DA and synthesis of gold nanoparticles and complexation of gadolinium ions on the HP backbone by FTIR spectroscopy. Attenuated total reflectance (ATR) FTIR spectra were recorded on a Thermo Fisher Scientific Nicolet 6700 FTIR in the infrared region (4000–650 cm$^{-1}$) with 32 scans at a resolution of 4 cm$^{-1}$.

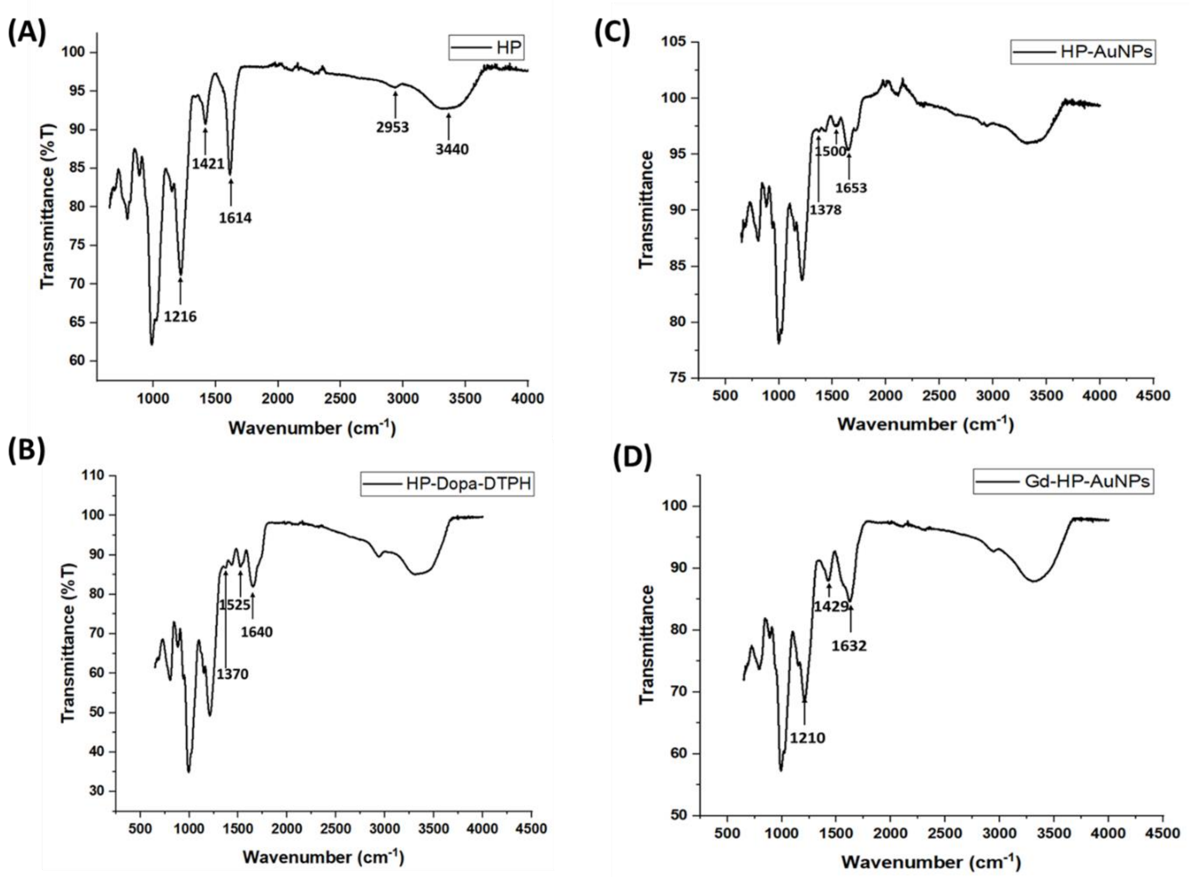


**Figure S5**: ATR-FTIR spectra of (A) heparin (HP), (B) conjugated dopamine and DTPH modified heparin (HA-Dopa-DTPH), (C) HP-AuNPs, and (D) Gd-complexed HP-AuNPs (Gd-HP-AuNPs)

#### *1.3 Synthesis of fluorescein tagged HP-AuNPs (FITC-HP-AuNP)*

The HP-AuNPs were tagged with the fluorescein molecule by thiosemicarbazide chemistry between available hydrazides on AuNPs and fluorescein isothiocyanate. Briefly, 60 mg HP-AuNPs (0.1 mmol) were dissolved in 51 mL de-ionized water. 4 mg fluorescein isothiocyanate (FITC, 0.01 mmol) was separately dissolved in 9 mL DMSO and dropwise added to the HP-AuNP solution. The reaction mixture was allowed to stir overnight. Thereafter, the reaction mixture was dialyzed using (Spectra Por-6, MWCO 3500 g/mol) against dilute HCl (pH = 3.5) containing 0.1 M NaCl (4×2L, 48 h), and then against deionized water (3×2L, 24 h). The solution was then lyophilized to obtain yellowish fluffy FITC-HP-AuNP conjugates. Finally, the material was washed with anhydrous MeOH to remove any traces of trapped free fluorescein isothiocyanate. The hydrodynamic size distribution was determined to be 127.7 nm with PDI 0.35 using a Zetasizer Nano ZS (Malvern, UK), while the surface zeta potential was -40.50±1.08 mV.

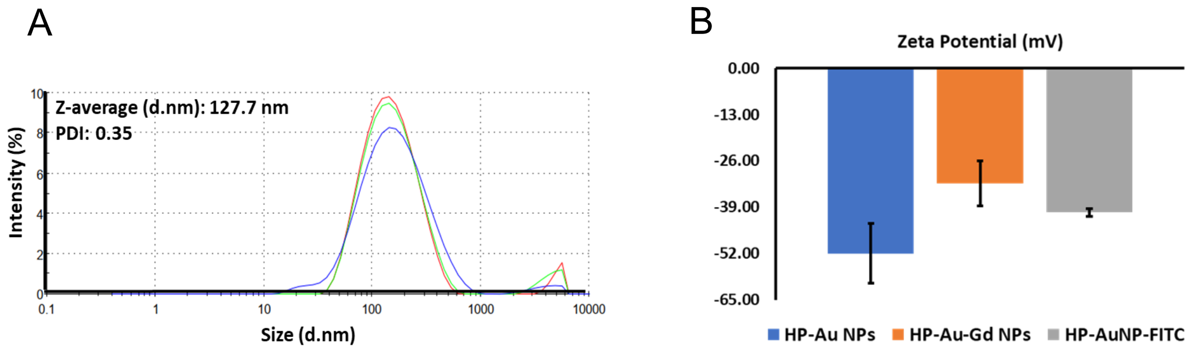


**Figure S6:** (A) Particle size distribution of FITC-HP-AuNPs in water (0.1 mg $ml^{-1}$) as determined by dynamic light scattering measurements. (B) Comparison of Zeta potentials of between HP-AuNPs (-51.97±8.35 mV), Gd-HP-AuNPs (-32.37±6.27 mV), and FITC-conjugated HP-AuNPs (-40.50±1.08 mV) (n=3).

#### *1.4 Radiolabeling of HP-AuNPs*

$^{68}$Ga ($t_{1/2}$ = 68 min, β+ = 89% and EC=11%) was available from a $^{68}$Ge/$^{68}$Ga-generator (Cyclotron Co., Obninsk, Russia) where the $^{68}$Ge ($t_{1/2}$ = 270.8 d) was fixed to a sorbent matrix based on modified titanium dioxide. The nominal $^{68}$Ge activity loaded onto the generator column was 1850 MBq (50 mCi). The $^{68}$Ga was eluted with 5 mL of metal-free 0.1 M hydrochloric acid (prepared from Ultra Trace Elemental Analysis Grade water, #W9-1, Fisher Chemical, and concentrated hydrochloric acid VWR Chemicals Ultrapure Normatom for trace metal analysis) using a syringe driver. The elution profile was determined by fractionating and measuring the $^{68}$Ga activity in each successive fraction of the eluate. The pH of the high activity $^{68}$Ge/$^{68}$Ga-generator eluate (600 µL, 224 MBq) used for the reaction was adjusted to pH ~ 4.0 by adding sodium acetate (140 µL, 1.0 M, pH 4.0) and NaOH (60 µL, 1.0 M). Then 2.0 mg HP-Au-Gd was added as a solution in 200 µL 0.10 M NaOAc (pH 4.0). The reaction mixture was heated to 70 °C in the heating block for 15 min.

##### *1.4.1 Purification of $^{68}$Ga-HP-AuNPs*

The reaction mixture (1000 µL) was purified in parallel using two NAP 5 columns (GE Healthcare, cutoff 5 kDa). First, the columns were equilibrated with 2 mL PBS each, then the reaction mixture (2×500 µL) was loaded. Each column was fractionally eluted with PBS (250, 750, and 1000 µL fractions, see Table 1). The load volume (500 µL) and first 250 µL contained no radioactivity and were discarded. The subsequent strongly purple-colored 750 µL fraction containing the majority of the $^{68}$Ga-HP-AuNPs was collected. Finally, the low molecular weight (LMW) fraction was eluted with 1000 µL. The fractions containing $^{68}$Ga-HP-AuNPs were pooled (92 MBq in 2×750 µL PBS, 46% RCY (decay corrected) calculated from the reaction mixture, 18% RCY (decay corrected) from a total eluted activity from the generator, in approx. 25 min after eluting the generator) before delivery for biological testing.

**Table 1.** Purification of $^{68}$Ga-HP-AuNPs

| | Radioactivity (MBq) | % RCY (decay corrected) |
|---|---|---|
| Reaction mixture | 224 | 100 |
| Left in reactor | 0.6 | 0.3 |
| Load volume 500 µL | 0 | 0 |
| Fr 1 250 µL (void) | 0 | 0 |
| Fr 2 750 µL ($^{68}$Ga-HP-Au) | 92 | 46 |
| Fr 3 1000 µL (LMW fraction) | 78 | 42 |
| Left in NAP5 | 20.8 | 11 |
| Total | | **99+ (recovery)** |

#### *1.4.2 Preparation of $^{68}$Ga-Acetate*

As a control, we prepared $^{68}$Ga-Acetate for in vivo experiments. $^{68}$Ga-Acetate should behave similarly to any $^{68}Ga^{3+}$ potentially released from $^{68}$Ga-HP-AuNP in vivo in the bloodstream. The high activity $^{68}$Ge/$^{68}$Ga-generator fraction (600 µL, 224 MBq) was first collected in a vial. The ratio of the reagents was adjusted for radioactive decay to give a final 40 MBq $^{68}$Ga-acetate in 500 µL at the time for injection. The synthesis was performed a maximum of 5 min prior to animal injection and separate $^{68}$Ga-acetate preparation was made for each animal.

For example, when the synthesis was started at 9:05, NaOH 1.0 M (11.5 µL, 11.5 µmol) was mixed with buffer NaOAc-buffer pH 4.6 (623 µL, Honeywell) in a 2 mL Eppendorf tube. $^{68}$Ga (60 MBq) in 0.10 M HCl (115 µL, 11.5 µmol) was added to a total volume of 750 µL and the mixture was vortexed. Immediately, the animal injection syringe was prepared by aspirating 0.50 mL (40 MBq).

#### *1.5 Synthesis of Gd-HP-AuNPs as MRI Contrast agents*

$Gd^{3+}$ ions were physically entrapped within the HP-AuNP system and stabilized by the available carboxylates and diol moieties of conjugated dopamine to the heparin backbone, yielding stable Gd-HP-AuNPs. To illustrate, 94 mg HP-AuNPs were dispersed in 90 ml deionized water. The pH of the solution was adjusted to 11 using 1 N NaOH. 19 mg gadolinium chloride hexahydrate (20 wt% of HP-AuNPs) was separately dissolved in 4 ml deionized water and dropwise added to the previous solution and allowed to stir overnight. The solution was next dialyzed (membrane MWCO: 3.5 KDa) against distilled water for 24 hours to remove any unreacted metal ions from the system. Then it was lyophilized to obtain Gd-HP-AuNPs as violet fluffy powder.  The hydrodynamic particle size distribution of Gd-HP-AuNP was determined to be 75.74 nm with PDI 0.257 using a Zetasizer Nano ZS (Malvern, UK) while the surface zeta potential was -32.37±6.27 mV.

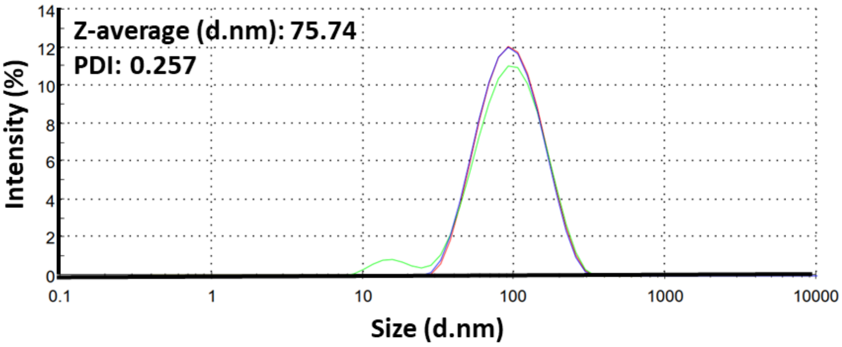


**Figure S7:** Particle size distribution of Gd-HP-AuNPs in water (0.1 mg $ml^{-1}$) as determined by dynamic light scattering measurements.

#### *1.5.1 Determination of Au and Gd amount by ICP-MS*

Gd and Au concentrations were measured by inductively coupled plasma mass spectrometry (ICP-MS) using Thermo Scientific iCAP™ RQ equipment. The instrument was equipped with a MicroMist borosilicate nebulizer 400 µl/min (Glass Expansion, Australia), a Peltier-cooled quartz spray chamber operating at 3 °C, a 2.5 mm ID quartz injector, a demountable quartz torch, a Ni skimmer cone, and a high matrix interface skimmer cone insert. All measurements were performed in Kinetic Energy Discrimination (KED) mode using He as collision gas in the collision/reaction cell and Ar as the carrier gas. The instrument was tuned using a solution containing 1 µg/l of Ba, Bi, Ce, Co, In, Li, and U for high sensitivity and minimum oxide levels (CeO/Ce < 2%). The measurement and analysis were performed using eQuant evaluation mode within Qtegra ISDS Software. Internal standard (1 µg/L U, 1 µg/L Rh, and 5 µg/L Ge) was added to the samples for the correction of nonspectral interferences in the analysis.

Ionic standard solutions with concentration range of 0.001–10 µg/L for Gd and Au were prepared in 2% aqua regia (35% w/v HCl: 65% w/v $HNO_3$; 4:1 v/v) + 1% thiourea and applied to measure the calibration curves. Thiourea was used in all the solutions to eliminate the Au memory effect in ICP-MS measurement.[4] Thus, all the measurements were made with 2 % aqua regia + 1% thiourea in the matrix. 197Au (KED) signal intensity < 500 cps (< 4 ng/L) was used as a wash-out criterion before the measurement of a new sample. The limit of detection (LOD) for Au was 2 ng/L. Metal-free polypropylene centrifuge tubes (VWR®), ultrapure deionized $H_2O$ (18.2 MΩ cm, Merck Milli-Q®), thiourea (>99.0%, ReagentPlus®, Sigma-Aldrich), mono-component reference solutions (Romil PrimAg®), and super pure acids (Romil-SpA™) were used in the sample preparation. Solid samples were wet digested in open vessels and diluted for the ICP-MS measurement.

Particle samples (m = 0.16–1.66 mg) were first dissolved in 1 ml $H_2O$. Then, 50 µL of sample solution was digested in 1 mL freshly prepared aqua regia at room temperature and finally diluted with 1% thiourea solution to obtain a final volume of 50 mL with 2% aqua regia concentration.

| Sample | Concentration (mg/ml) | Dilution factor | Au (wt%) | Gd (wt%) |
|---|---|---|---|---|
| **Gd-HP-AuNPs** | **1** | **1000** | **3.51** | **8** |

**Table S1:** Weight percentage of Au and Gd in HP-Au-Gd complex as determined by ICP-MS

#### *1.5.2 Thermogravimetric analysis (TGA)*

The thermogravimetric analysis was further performed to calculate the polymer contents in the synthesized HP-AuNPs and Gd-HP-AuNPs using an $Al_2O_3$ pan and the temperature was increased to 1000 °C at 20 °C/min under the $N_2$ flow.

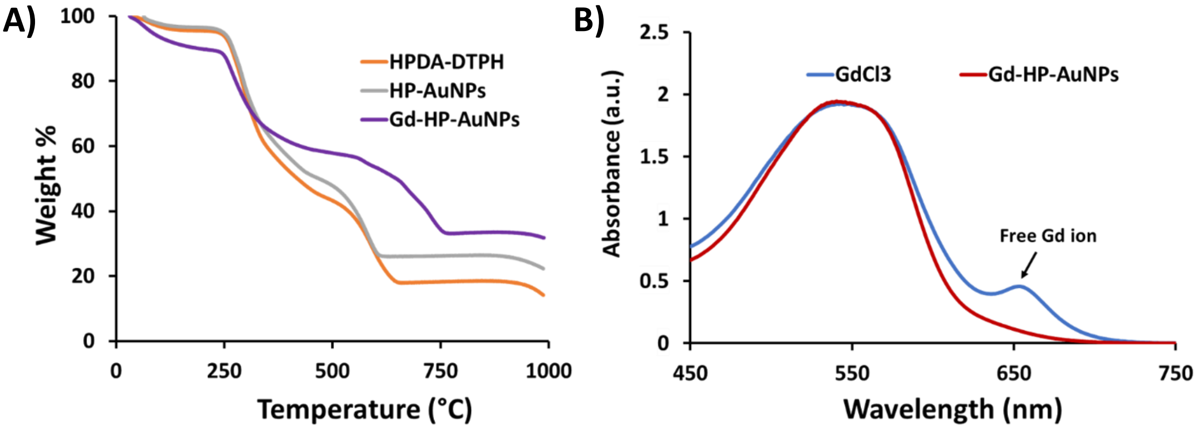


**Figure S8:** (A) Thermogravimetric analysis of conjugated dopamine and DTPH modified heparin (HA-Dopa-DTPH), HP-AuNPs, and Gd-complexed HP-AuNPs (Gd-HP-AuNPs), and (B) Determination of free Gd ion by Arsenazo (III) assay

#### *1.5.3 Free Gd Determination by Arsenazo III colorimetric assay*

Free $Gd^{3+}$ was determined to be 1.7 weight% by the absorption of Gd-Arsenazo III complex following the published protocol.[5] Briefly, equal molar ratio of arsenazo III solution was mixed with of free $Gd^{3+}$, and then the absorbance spectra were measured using with a UV-vis spectrometer (V-570 UV/VIS/NIR Spectrophotometer, JASCO, Japan) at 660 nm to obtain the standard curve. Finally, HP-Au-Gd was dissolved in 1x PBS (pH 7.4) at 1 mg/mL and mixed with arsenazo III solution at a 1:1 volume ratio. The spectra were recorded at 660 nm and the free $Gd^{3+}$ percentage was determined from the standard curve.

#### *1.5.4 Longitudinal Relaxivity ($r_1$) Measurement*

The longitudinal relaxation time $T_1$ of Gd-HP-AuNPs at different Gd concentrations was evaluated at room temperature by using a 1.0 T-MRI scanner (ICON, Bruker Biospin, Ettlingen, Germany). The contrast agents were initially put into syringes and set in the center of the volume coil. The sample temperature was maintained at room temperature. By using the MRI scanner, horizontal single-slice T1-weighted MR images were acquired (spin echo, TR/TE = 400/10 ms, slice thickness = 2.0 mm, matrix = 256 × 256, field of view (FOV) = 38.4 × 38.4 mm, number of averages (NA) = 10, number of slices = 1). Horizontal single-slice inversion-recovery MRI was done to calculate the $T_1$ and $r_1$ by using rapid acquisition with relaxation enhancement (RARE) acquisition (TR = 20000 ms, TE = 17 ms, inversion time =

45, 100, 200, 400, 800, 1600, 3200, 6400, 8000, 10000, 12000 ms, matrix size = 64 × 64, FOV = 38.4 × 38.4 mm, slice thickness = 2.0 mm, RARE factor = 4, and NA = 1). Then, the $r_1$ relaxivities were calculated by using the following equation:

$r_1 = (1/T_1 - 1/T_1(0)) / [Gd]$

where, [Gd] is the concentration of Gd, and $1/T_1(0)$ and $1/T_1$ are the longitudinal relaxation rate without and with paramagnetic species, respectively.

### 2. In-vitro assays

#### 2.1 Evaluation of nanoparticle (HP-FTSC and CS-FTSC) permeability across an in vitro BBB model

##### 2.1.1 *Cell culture*

All the cells were maintained and grown at 37 ºC in a humidified atmosphere containing 5 % $CO_2$ unless stated otherwise for not more than 10 and 15 passages. Mycoplasma contamination was routinely checked using the mycoplasma detection kit (11-1050, Minerva Labs).
Immortalized Human Umbilical vein endothelial cells (HuAR2T) were expanded in EBM-2 medium (c-22211, Promocell) with SupplementMix (c-39216) and 2 µg/ml doxycycline. Murine brain microvessel endothelial cells (bEND3) were expanded in DMEM (21331-020, Gibco) containing 1g/l glucose, and with 10% FBS, 1% L-glutamine, 100 U/ml penicillin, and 100 µg/ml streptomycin, Normal Human Astrocytes (NHA, CC-2565, Lonza) were maintained in ABM (CC-3187) with SingleQuots supplements (CC-4123). HIFko Mouse astrocytes were expanded in BME-1 (21010046, ThermoFisher) supplemented with 5% FBS, 5 mL 1 M HEPES, mL 100 mM sodium pyruvate, 3g glucose, and 5 mL penicillin/streptomycin.

##### *2.1.2 In vitro Blood-Brain Barrier penetration*

Murine and Human blood-brain barriers (BBB) in a dish were established according to a previously published protocol.[6] Briefly, immortalized mouse brain microvessel endothelial cells (bEND3) or human immortalized umbilical vein endothelial cells (HuAR2t) were co-cultured in serum-free conditions in Transwell inserts with immortalized mouse astrocytes (HIFko) or Normal Human Astrocytes (NHA), respectively. After 5 to 7 days of co-culture required for the formation of a tight impermeable endothelial cell layer, the BBB permeability is determined by measuring the small molecular weight probe sodium fluorescein diffusion from the blood to the brain side. The following formula is used for the permeability calculation:

Permeability = $\frac{dQ}{dT \times A \times C^0}$

Expected permeability values for the murine and humanized BBB are in the $10^{-6}$ and $10^{-5}$ cm/s range, respectively.

To evaluate the nanoparticles transportation through the BBB, 100 µg of fluorescently labeled HP and CS polymers (HP-FTSC and CS-FTSC) were added to the endothelial side of the inserts. After 48h, samples from both the blood and brain sides were collected and nanoparticle-related fluorescence was measured on a fluorescence plate reader to determine the permeability of the BBB for each construct. BBB inserts were then fixed on ice-cold 4% PFA (10 min) and nuclear counterstaining with DAPI (1 µg/mL, Sigma). The stained membranes were eventually mounted on microscope slides with antifading Mowiol 4-88 solution (Sigma-Aldrich) and imaged on a Zeiss LSM880 confocal microscope.

#### *2.1.3 MTT- proliferation assay*

5000 cells/well were plated on 96-well plates in 3-10 replicates (100 µL volume). At the indicated time points, 10 µL of 3-(4,5-Dimethylthiazol-2-yl)-2,5-Diphenyltetrazolium Bromide (MTT) (5 mg/ml in PBS) was added, and the cells were incubated for 2 hours. Finally, the cells were lysed (10% SDS, 10 mM HCl) o/n and the absorbance was measured at 540 nm using Multiskan Ascent software version 2.6 (Thermo Labsystems).

### 2.2 Mechanism of the passage through the BBB

#### *2.2.1 In-vitro gene expression analysis of gap junction proteins*

To analyze the expression of gap junction proteins, we cultured 60,000 murine microvascular brain endothelial cells (bEND3) in a 24-well plate that were incubated overnight at 37˚C, 5% $CO_2$. Thereafter, the medium (DMEM (Gibco) containing 10% fetal bovine serum (Gibco, South American) and 1% penicillin-streptomycin (Gibco)) was changed and 500 µL of 0.2 mg/mL HP-AuNP-FITC suspended in serum-free medium was added to cells and were incubated for 24h. Untreated cells were used as control. After 24 h, RNA was extracted using the RNeasy Plus Mini kit from Qiagen. 300 ng of the total RNA was used to make the cDNA. The cDNA was prepared using High capacity RNA to cDNA (Thermo Fisher) kit according to the manufacturer's protocol and qRT-PCR was performed with cDNA and TaqMan® Fast Advanced Master Mix (2X) (Applied Biosystems). The real-time PCR reactions were carried out with 10 µl of 2x TaqMan® Universal PCR Master Mix, 2 µl cDNA, and 1 µl of TaqMan gene-specific assay mix (Transferrin Receptor, Tight junction protein 1 (ZO1), Claudin 5 (CLDN5), Glucose transporter 1 (Glut1) and β-actin) (Applied Biosystems) in a 20 µl final reaction volume. Reference gene, β-actin (ACTB) (Taqman primers, Thermo Fisher) were selected as a control for normalization of real-time PCR data. The amplification was carried out using the Biorad CFX1000 (Bio-rad) using a 40-cycle program. The CFX manager software automatically calculates the raw Ct (cycle threshold) values. Samples were normalized relative to endogenous control and differences in cycle number thresholds were calculated using the comparative quantitation 2−ΔΔCT method.

**A**

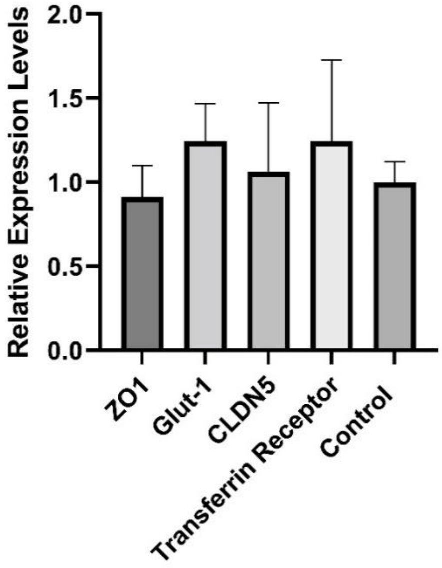


**B**

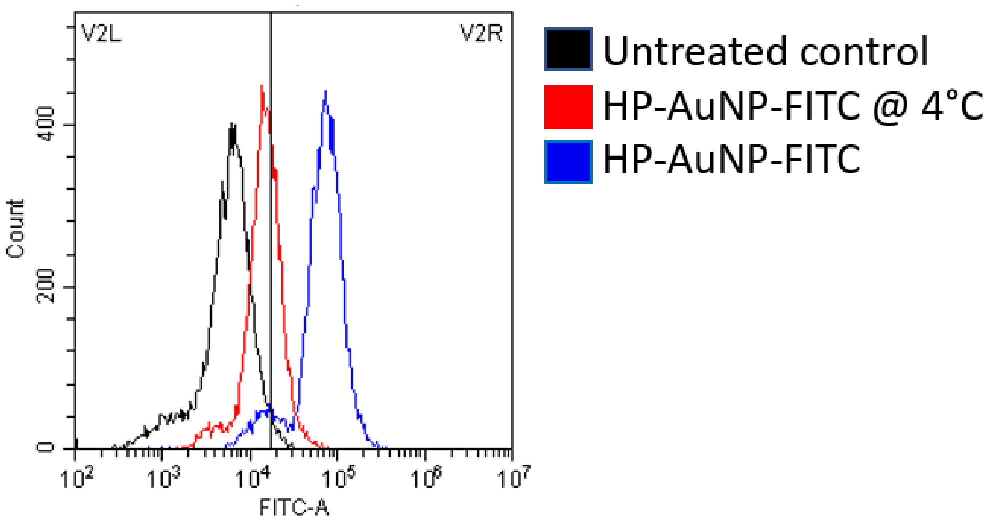

**Figure S9 (A):** The expression levels of the gap junction protein ZO1, CLDN5, Glut1, and transferrin receptor. (**B**) Flow cytometry histogram showing reduced uptake of HP-AuNP-FITC by microvascular murine brain endothelial cells (bEND3 cells) at 4°C.

#### *2.2.2 In-vitro HP-AuNP-FITC uptake and blocking studies*

For the uptake study, 50,000 murine microvascular brain endothelial cells (bEND3) were plated in a 24-well plate and incubated overnight at 37˚C, 5% $CO_2$. Thereafter, the medium (DMEM (Gibco) containing 10% fetal bovine serum (Gibco, South American) and 1% penicillin-streptomycin (Gibco)) was changed and 500 µL of 0.1 mg/mL HP-AuNP-FITC suspended in serum-free medium was added to cells and were incubated for 2h. The cells were then washed 3 times with PBS and trypsinized to form a single-cell suspension and were subsequently used for flow cytometry using a Cytoflex (Beckman Coulter) flow cytometer. For the blocking study, the cells were incubated at 4°C for 1h after which the HP-AuNP-FITC at 0.1mg/mL and incubated for 2 h. The cells were then washed with PBS prior to trypsinization, and the same steps were followed as presented before.

### 3. In-Vivo assays

#### 3.1 Comparative HP-FTSC and CS-FTSC diffusion across the BBB in adult male FVB mice

All experiments involving animals were authorized by the National Animal Experimental Board in Finland (Helsinki, Finland), under the licenses ESAVI/6285/04.10.07/2014 and ESAVI/403/2019. Prior to the infusion, stock lyophilized HP-FTSC and CS-FTSC stocks were reconstituted in ice-cold PBS and briefly homogenized in a water bath sonicator and filtered using a 0.22 µm syringe filter. Three male FVB/NRj (Lab Animal Center, Helsinki, Finland) mice (n = 3) were injected with 4 mg/kg (100 µg in 200 µl PBS) of HP-FTSC or CS-FTSC or vehicle solution through the caudal vein. After 1 h or 24 h post-injection, the mouse was deeply anesthetized with a ketamine/xylazine mix and perfused intracardially with ice-cold PBS, and brains were collected and snap-frozen in -50°C isopentane (Honeywell) upon tissue processing and nanoparticle-related fluorescence quantification as described below.

Snap-frozen xenografted brains were cut using a cryotome (Cryostar NX70, Thermo Scientific) into a series of 30 µm thick mouse brain sections. Tissue sections were fixed in 4% PFA, blocked with 5% FBS and 0.03% Triton X (Sigma). The frozen tissue sections were stained by immunofluorescence for podocalyxin (PODXL, R&D Systems MAB1556) that show the blood vessels and DAPI that stain the nucleus. Brain sections were then imaged on a confocal microscope (Zeiss LSM 880).

### 3.2 Biodistribution of HP-AuNP-FITC in healthy C57BL/6JRj, FVB/NRj, and NMRI-Nude mice

#### *3.2.1 Animal experiments*

Prior to the animal experiment, the lyophilized nanoparticle stocks were reconstituted in ice-cold PBS, briefly homogenized in a water bath sonicator, and filtered using a 0.22 µm syringe filter. To evaluate the brain biodistribution and cell-specific uptake of HP-AuNPs, 8 to 12 weeks-old female C57/Bl6 nrj (n=5), female immunocompromised NMRI-Nude mice (n = 5) (Janvier Labs) and male FVB/NRj (Lab Animal Center, Helsinki, Finland) mice (n = 5) were injected with 4 mg/kg (100 µg in 200 µl PBS) HP-AuNP-FITC or vehicle solution through the caudal vein. Animals were then deeply anesthetized with a ketamine/xylazine mix at 3 h post-injection, the animals were perfused intracardially with ice-cold PBS, and brains were collected and snap-frozen in -50°C isopentane (Honeywell). The tissue samples were processed, and the nanoparticle-related fluorescence quantification was performed as described below.

#### *3.2.2 Animal tissue processing and analysis*

Snap-frozen xenografted brains were cut using a cryotome (Cryostar NX70, Thermo Scientific) into a series of 9 µm-thick coronal sections collected from the frontal to the anteroposterior part of the diencephalon. Tissue sections were fixed in 4% PFA, blocked with 5% FBS and 0,03% Triton X (Sigma), and stained by immunofluorescence for neuron cell adhesion molecule (NCAM1, Abcam ab75813), neuropilin-1 (NRP-1, Cell Signalling #3725), podocalyxin (PODXL, R&D Systems MAB1556), glial fibrillary acidic protein (GFAP, Abcam ab53554), allograft inflammatory factor (IBA-1, Abcam ab107159) or with an antibody directed against FITC (R&D Systems G-148-C). Brain sections were then imaged on a confocal microscope (Zeiss LSM 880). Quantification of the different nanoparticle distribution features (location into the brain, association with a specific brain cell type) was performed on 10 sections equally distributed along the whole diencephalon using the CellProfiler (https://cellprofiler.org/) and ImageJ (https://imagej.nih.gov/ij) software.

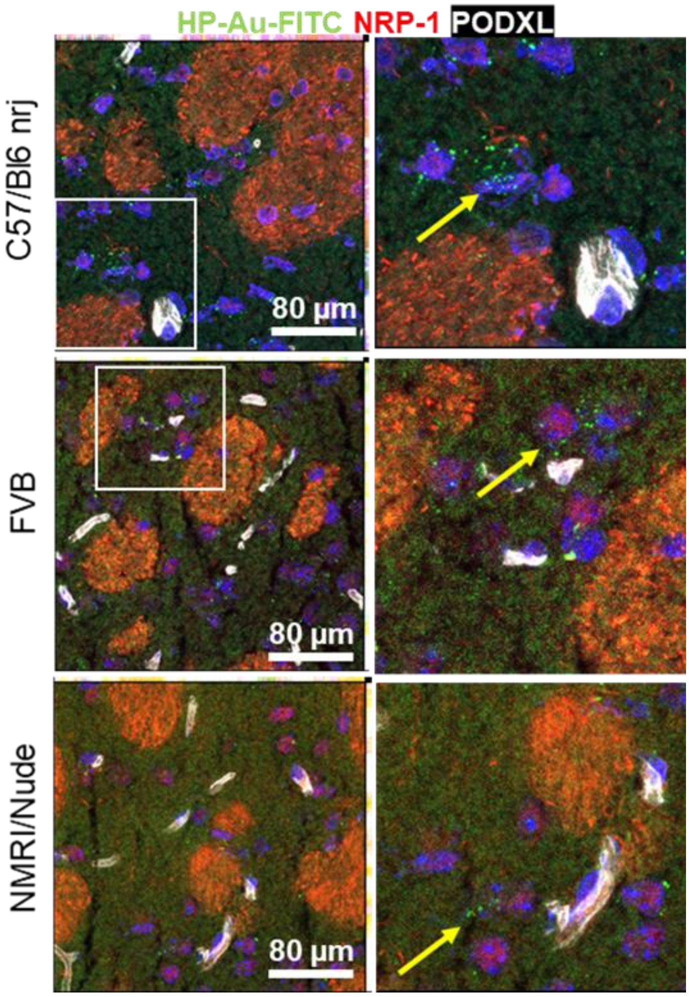


**Figure S10:** Brain micrographs of female C57/Bl nrj, NMRI-Nude, and male C57/Bl6 mice injected intravenously with 100 µg of FITC-HP-AuNPs (green) for 3 h. Pack of axons/nerves from the striatum were labeled with an NRP-1 antibody (red) and brain endothelial cells with a PODXL antibody (white). Arrows are pointing toward the classical distribution of the nanoparticles, e.g., found isolated in the brain parenchyma.

### 3.2 Biodistribution study of $^{68}$Ga-HP-AuNPs in Sprague Dawley rats by PET/MRI

#### *3.2.1 Biodistribution of $^{68}$Ga-HP-AuNPs*

Biodistribution of $^{68}$Ga-HP-Au and 68Ga-Acetate was evaluated in Sprague Dawley rats (n = 6 for each radiotracer, male, healthy, 297.0±18.9 g) by in vivo PET/MRI, plasma stability, and an ex vivo autoradiography binding assay.

All procedures involving animals were in compliance with the ARRIVE guidelines, approved by the Animal Ethics Committee of the Swedish Animal Welfare Agency, and carried out in accordance with the relevant national and institutional guidelines ("Uppsala university guidelines on animal experimentation", UFV 2007/724).

Anesthetized animals were administered 23.0±5.0 MBq (corresponding to 500µg HP-Au) $^{68}$Ga-HP-Au (n=6) or 39.6±2.5 MBq 68Ga-Acetate (n=6) via the tail vein. Three of the rats of each group were used for PET/MRI imaging and the others for ex vivo autoradiography studies, and plasma stability.

#### *3.2.2 PET/MRI imaging and analysis*

The rats used for in vivo distribution (n=3 from each group) were examined by whole-body PET/MRI. Dynamic PET images were acquired from injection to 90 min using a small animal PET-MRI system (nanoPET/MRI, 3T, Mediso, Hungary). Dynamic whole-body PET scanning was performed using multiple whole-body sweeps (3 beds per pass: 1 x 1 min, 3 × 3 min, 2 × 20 min, 2 × 30 min). After the PET acquisition, the animals were euthanized by sodium thiopental (Apoteket AB, Stockholm, Sweden) while lying in the scanner. MRI sequences were realized post-mortem for 30 min (Spin-Echo Multi-FOV sequence). PET images were reconstructed by the use of the Maximum Likelihood Estimation Maximized (MLEM) algorithm (10 iterations). PET images were analyzed in PMOD 4.0 (PMOD Technologies, Zürich, Switzerland). Briefly, tissues of interest (brain, heart ventricle, liver, spleen, bladder) were segmented on PET summation images and co-registered MRI images. PET uptake was readout for all time points, decay corrected, and expressed as Standardized Uptake Values (SUV) or Percentage of the injected dose per gram of tissue (%ID/g). The total uptake in each organ (%ID) was estimated by multiplying the tissue concentration (%ID/g) with the volume of the entire organ (g).

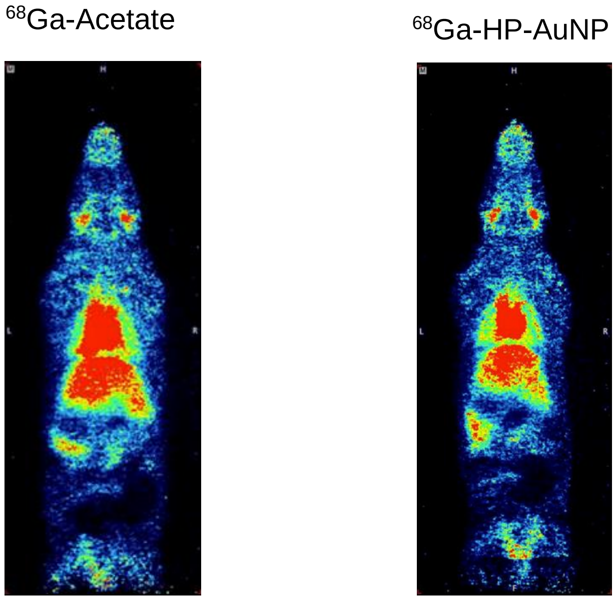

**Figure S11:** Dynamic whole-body imaging of administered $^{68}$Ga-Acetate and $^{68}$Ga-HP-AuNP in Sprague Dawley rats (the video file is attached separately in PPT)

#### *3.2.3 Plasma stability of $^{68}$Ga-HP-Au and $^{68}$Ga-Acetate*

Plasma stability was evaluated in n=3 rats from each group. Blood was withdrawn from the tail vein to heparinized Eppendorf tubes at 5-, 30- and 60-minutes post radiotracer administration and immediately cooled on ice. A small aliquot (approx. 50 µL) of whole blood was weighed and measured for radioactivity in a well counter. The remaining blood was centrifuged (4000 RCF, 4 °C, 5 min) to separate RBC and plasma. Aliquots of plasma (100 µL) and RBC (100 µL) were weighed and measured for radioactivity.

The radioactivity was decay corrected to the time point of injection and normalized to the amount of injected radioactivity to achieve the decay corrected %ID/g. There was no normalization for the different bodyweight of the animals.

The 100 µL plasma sample was diluted with cold PBS (1400 µL) and then centrifuged in a centrifugal filter (Amicon Ultra 2 mL Ultracel® 10k regenerated cellulose 10 000 NMWL, 4 °C) until the liquid surface reached the lower rim of the filter. Additional 700 µL PBS was added and centrifuged in a similar way once more. The activity in the filtrate was measured which corresponds to LMW unbound $^{68}$Ga while the remaining activity in the filters represents all HMW compounds including intact $^{68}$Ga-NP.

When 0.5 mL Amicon filters were used, the amount of plasma was decreased to 25 µL and diluted with PBS (475 µL) and then rinsed with PBS (200 µL). However, this resulted in significantly lower HMW-fraction compared to the 2.0 mL filters. Hence the results were omitted in figure 4. All radioactivity data is decay corrected to the time-point for injection and normalized for the sample weight and the injected amount.

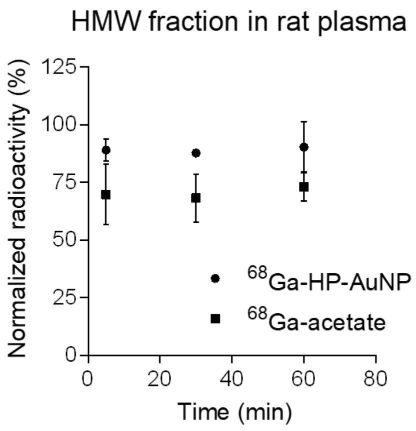

**Figure S12:** Normalized radioactivity in rat blood plasma after tail vein infusion of $^{68}$Ga-HP-AuNPs and $^{68}$Ga-acetate from 7 min. to 1 h post-injection

#### *3.2.4 Ex vivo autoradiography binding assay*

After the rats used for plasma stability assessment were euthanized 60 minutes after radiotracer administration, the liver, spleen, and brain were immediately collected for freezing in isopentane chilled with dry ice. The frozen tissue samples were sectioned to 20 μm sections with a cryostat microtome (Micron HM560, Germany) and mounted on Menzel Super Frost plus glass slides. The sections were exposed to phosphorimaging plates for 3h and scanned by a Phosphorimager system (Amersham Typhoon IP, GE Health). The sections were visualized and analyzed using the software ImageJ (ImageJ 1.45S, NIH, Bethesda, USA). The same sections were then stained with hematoxylin for histological analysis.

### 3.3 Evaluation of Gd-HP-AuNPs localization in female BALB/c mice in a Xenograft model

#### *In vivo MRI imaging*

Female BALB/c mice (6-weeks-old) were inoculated intracranially with U87-MG cells (1 × 104 cells). After 15 days, the animals were studied by MRI after intravenous injection of Gd-HP-Au NPs at a dose of 10 mg/kg based on Gd. The MRI measurements were done in 1T MRI equipment (Bruker BioSpin) with the following imaging parameters: spin-echo method, TR = 400 ms, TE = 12 ms, FOV = 30 × 30 mm, matrix size = 188 × 188, and slice thickness = 1.8 mm. Images were acquired at defined time points.

(3) Lee, K.; Lim, S.; Go, N.; Kim, J.; Mun, J.; Kim, T. H. Dopamine-Grafted Heparin as an Additive to the Commercialized Carboxymethyl Cellulose/Styrene-Butadiene Rubber Binder for Practical Use of SiOx/Graphite Composite Anode. *Sci. Reports*

*2018 81* **2018**, *8* (1), 1–9. https://doi.org/10.1038/s41598-018-29705-y.

(4) Chen, W.; Wee, P.; Brindle, I. D. Elimination of the Memory Effects of Gold, Mercury and Silver in Inductively Coupled Plasma Atomic Emission Spectroscopy. *J. Anal. At. Spectrom.* **2000**, *15* (4), 409–413. https://doi.org/10.1039/A908658G.

(5) Mi, P.; Cabral, H.; Kokuryo, D.; Rafi, M.; Terada, Y.; Aoki, I.; Saga, T.; Takehiko, I.; Nishiyama, N.; Kataoka, K. Gd-DTPA-Loaded Polymer–Metal Complex Micelles with High Relaxivity for MR Cancer Imaging. *Biomaterials* **2013**, *34* (2), 492–500. https://doi.org/10.1016/J.BIOMATERIALS.2012.09.030.

(6) Le Joncour, V.; Karaman, S.; Laakkonen, P. M. Predicting in Vivo Payloads Delivery Using a Blood-Brain Tumor-Barrier in a Dish. *J. Vis. Exp.* **2019**, *2019* (146). https://doi.org/10.3791/59384.
